# Deciphering the Therapeutic Potential of Resveratrol Against Pancreatic Cancer Through Network Pharmacology

**DOI:** 10.64898/2026.01.27.701943

**Authors:** Aditya Bisen, Roshini Singh, Varun Jaiswal, Sanjay Mishra, Varun Kumar Shrama, Manoj Kumar Mishra

## Abstract

Resveratrol, a naturally occurring polyphenol, exhibits anticancer, anti-inflammatory, and antioxidant properties. However, its molecular mechanisms in pancreatic cancer remain incompletely understood, necessitating integrative approaches such as network pharmacology and molecular docking. Potential resveratrol targets were identified using Swiss Target Prediction, while pancreatic cancer–related genes were retrieved from GeneCards and NCBI Gene databases. Overlapping targets were obtained through Venn analysis, followed by protein– protein interaction network construction, hub gene selection, and enrichment analysis. Molecular docking validated compound–target interactions. A total of 100 predicted resveratrol targets and 1,447 pancreatic cancer– associated genes were screened, yielding 39 overlapping genes. Network analysis identified hub genes including EGFR, SRC, MTOR, PIK3CA, PIK3CB, BCL2, and PTGS2. Gene Ontology enrichment indicated roles in cell proliferation, apoptosis regulation, inflammatory response and metabolic regulation while KEGG pathway analysis highlighted the PI3K–Akt, ErbB, and EGFR inhibitor resistance signaling as being closely associated with pancreatic cancer pathway. Docking analysis revealed strong binding of resveratrol with KRAS (–8.2 kcal/mol), EGFR (–7.9 kcal/mol), and MTOR (–7.7 kcal/mol), stabilized by hydrogen bonding. The interaction with KRAS, although not among the predicted targets. Expression profiling validated upregulation of hub genes in tumor samples. Resveratrol exerts multi-targeted effects in pancreatic cancer by modulating oncogenic pathways, particularly KRAS and PI3K/Akt/mTOR signaling. Its favourable safety profile and robust hub gene interactions highlight its potential as a supplementary therapeutic drug, necessitating further preclinical and clinical confirmation.

## Introduction

Pancreatic cancer is a growing global health concern, with over 510,000 new cases and 467,000 deaths reported in 2022, making it the sixth leading cause of cancer-related deaths (Leiphrakpam et al. 2025). Its incidence is projected to nearly double by 2050, primarily due to an aging global population and improved access to healthcare services (Zottl et al. 2025). Despite some progress, the five-year survival rate remains low at around 10% due to late diagnosis and lack of effective screening. Pancreatic ductal adenocarcinoma (PDAC) is the most common subtype, and the disease’s non-specific symptoms and silent progression highlights the urgent need for improved early detection and a deeper understanding of its molecular mechanism (Wong et al. 2023). The poor survival rate in pancreatic cancer is largely due to late diagnosis and limited treatment efficacy. Risk factors such as smoking, alcohol use, inactivity, and unhealthy diets contribute to disease progression (Jamal and Khan 2025). Molecular differences in tumors emphasize the need for personalized therapies. While chemotherapy and targeted treatments like Folfirinox, Erlotinib, and Olaparib are commonly used, their effectiveness is limited and often have severe side effects (Costa et al. 2015. Min and Lee 2022). As a result, there is growing interest in plant-based therapeutics, due to their multiple biological targets and reduced toxicity. Independent systems of traditional medicine, such as Ayurveda (India), Kampo (Japan), Sa-sang (Korea), and Traditional Chinese Medicine (China), have long utilized plants for healing purposes (Yuan et al. 2016). Globally, around 70,000 plant species have been used in traditional healthcare, with more than 3,000 species reported to exhibit potential anticancer properties. Notably, over 60% of current chemotherapeutic agents are derived from plants, including paclitaxel, vincristine, vinblastine, etoposide, irinotecan, and topotecan (Sulaiman et al. 2025). The integration of herbal medicines with modern cancer treatment represents a growing trend in integrative oncology, where herbal extracts are applied as supportive therapies to reduce side effects, enhance efficacy, and improve patient well-being. Documenting and scientifically validating traditional knowledge is therefore critical, as it provides an invaluable resource for novel drug discovery and development in oncology (Imtiaz et al. 2024).

Resveratrol, a naturally occurring stilbene polyphenol predominantly found in *Polygonum cuspidatum* and various dietary sources such as grapes, berries, and peanuts, has attracted considerable attention for its wide range of biological activities such as anticancer, anti-inflammatory, and antioxidant properties (Lee et al. 2014, Wu et al. 2018). Preclinical studies have shown its effectiveness in inhibiting tumor growth, inducing apoptosis, and regulating key cancer-related pathways (Salla et al. 2024). However, its clinical application is hindered by low bioavailability, prompting research into nanoparticle-based delivery systems. Mechanistically, resveratrol acts through DNA degradation via copper ions, downregulation of SphK2, and epigenetic modulation, showcasing its multi-targeted potential against complex diseases like pancreatic cancer (Daina et al. 2017). Despite these promising findings, the comprehensive molecular mechanisms underlying resveratrol’s effects in pancreatic cancer remain insufficiently understood. Most existing studies fail to fully describe its direct interactions with pancreatic cancer-specific targets. Moreover, the absence of integrative methodologies has constrained the exploration of its multi-target potential across the diverse Signaling networks implicated in pancreatic tumorigenesis. Network pharmacology, an emerging systems-level approach, integrates computational biology, genomics, and systems biology to systematically evaluate drug–target–disease interactions (Das et al. 2023). By leveraging big data, this approach not only facilitates the identification of synergistic effects in cancer therapy but also provides mechanistic insights into bioactive compounds.

In this study, we apply a network pharmacology approach to investigate the therapeutic potential of resveratrol in pancreatic cancer. This strategy enables the systematic identification of hub genes and pathways associated with its anticancer activity and provides a framework for clarifying its pleiotropic mechanisms. Ultimately, these insights may support the development of resveratrol as an adjunct or alternative therapeutic option for pancreatic cancer.

## MATERIALS AND METHODS

### ADMET Screening and Pharmacokinetics of Stilbene Compounds

Computational evaluation of physicochemical and pharmacokinetic properties was done using SwissADME (http://www.swissadme.ch/). Two important parameters, Lipinski’s rule and Veber’s rule, play a crucial role in evaluating the physicochemical properties of compounds. Investigating these properties is important for determining whether the compound is viable drug candidate or not. Additionally, ADMET properties such as Absorption, Distribution, Metabolism, Excretion, Toxicity, BBB (Blood brain barrier) permeability and GI (Gastrointestinal Tract) absorption were also evaluated (Pires et al. 2015). PkCSM (biosig.lab.uq.edu.au ) tool was used for evaluating Pharmacokinetic properties such as Skin Permeability, CYP2D6 Inhibitor, CYP3A4 Substrate, Herg Inhibitor 1 and 2, CNS (Central Nervous System), Oral Rat Acute Toxicity, Oral Rat Chronic Toxicity (Xu et al. 2015).

#### Toxicity Analysis of Resveratrol

Toxicity of RVT was predicted using the Pro-Tox-3 (https://tox.charite.de) web tool for finding toxicity activeness with respect to Hepatotoxicity, Neurotoxicity, Cytotoxicity, Nephrotoxicity, Carcinogenicity, Immunotoxicity, Mutagenicity (Banerjee et al. 2024).

#### Screening of Potential Targets genes of Resveratrol

The canonical SMILES of Resveratrol were obtained from PubChem Database (https://pubchem.ncbi.nlm.nih.gov/compound/44257189) and were given as an input for predicting Potential Targets of Resveratrol using Swiss Target Prediction (http://www.swisstargetprediction.ch/predict.php) (Daina et al. 2017, Kim et al. 2024).

### Acquisition of Gene Targets for Pancreatic Cancer

Potential gene targets associated with pancreatic cancer were identified using the Gene Cards database (https://www.genecards.org/) and with a relevance score ≥10 along with the NCBI Gene database where genes were sorted based on their high Relevance Score by applying keywords such as pancreatic cancer adenocarcinoma and pancreatic cancer (Stelzer et al. 2016, Wiegers et al. 2025). High-relevance genes were retrieved from GeneCards, while additional targets reported from previous experimental studies were collected through literature mining (Matsuoka 2016, Pergolizzi and Brower 2024). These literature-reported (wet-lab) genes were then compared with computationally predicted (dry-lab) targets to generate a comprehensive dataset for further analysis. This comparative strategy ensures that the computational predictions are cross validated with experimentally verified evidence, thereby enhancing the reliability and biological relevance of the identified gene set.

#### Identification of overlapping genes

A Venn diagram was constructed to identify and illustrate the genes that overlap between the resveratrol and pancreatic cancer. The intersected targets between the PC-related targets and resveratrol respectively were retained using Venny 2.1. (bioinfogp.cnb.csic.es) (Oliveros 2007, Agrawal et al. 2023).

### Protein-Protein Interaction (PPI) network construction of drug-potential targets

Protein-protein interaction (PPI) network construction was done using STRING database (https://string-db.org/), which contains data of known human genes. The common overlapping target gene were given as input in the database and employed to construct a PPI network model with the species set to “*Homo sapiens*”, and by setting the high confidence score at 0.700 (Szklarczyk et al. 2022).

### Identification of Hub Genes and construction of Compound-Target Interaction Network

After establishing the PPI network, the Cytoscape plug-in CytoHubba was used to filter out hub genes. Top 7 hub gene were selected by calculating the degree of each node in the PPI network. The Compound-Target Interaction Network was also constructed (Shannon et al. 2003. Chin et al. 2014).

#### Retrieving Gene Ontology (GO) and Hub Gene Pathways Data

DAVID database was used to obtain KEGG pathway analysis and Gene Ontology information (davidbioinformatics.nih.gov) by entering the Top 5 hub genes and selecting Homo Sapiens in species. The Gene Ontology comprised of Hub genes involved in various Biological Processes (BP), Cellular Component (CC) and Molecular Function (MF) along with Hub genes involved in various Signaling pathways along with their Gene count (Sherman et al. 2022).

### Enrichment Gene Ontology (GO) data using Shiny GO

The Shiny GO tool was used to construct Bar plots for various biological processes (BP), Cellular Component (CC), Molecular Function (MF) and KEGG pathways. The cutoff was set at 0.05 probability score to select top 25 BP, CC, MF and KEGG pathways and represent them by drawing conclusions from the bar plots created using ShinyGo version 0.82 (bioinformatics.sdstate.edu on 1 June 2025) (Ge et al. 2019).

#### Active site prediction

The active sites of the target protein were predicted using the CASTp 3.0 server (Computed Atlas of Surface Topography of Proteins) (Tian et al. 2018). The protein structure was analyzed to identify surface pockets and internal cavities, and pockets were ranked based on area and volume using the default probe radius of 1.4 Å. The top-ranked pocket and its associated amino acid residues were selected for defining the binding site in subsequent molecular docking studies.

#### Molecular docking studies

To validate protein-ligand complex interactions, molecular docking was performed using AutoDock Vina (Eberhardt et al. 2021). The structure of compounds was downloaded from PubChem and converted to PDBQT format using Open Babel. All the 3D structure of target protein were obtained from the RCSB Protein Data Bank (https://www.rcsb.org/). Proteins were prepared by removing water molecules, adding polar hydrogens, and calculating Gasteiger charges using AutoDock Tools (Eberhardt et al. 2021). Site-specific docking was carried out within a defined grid box around the active site, and the results were analysed based on binding affinity. The best docking poses were visualized using PyMOL and Discovery Studio Visualizer to assess binding interactions (Visualizer, D. 2005, Janson et al. 2016).

#### Validation of hub genes

The expression levels of the identified hub genes were validated using the GEPIA2 web server (http://gepia.cancer-pku.cn/). mRNA expression profiles in pancreatic cancer tissues were compared with normal controls based on TCGA and GTEx datasets. Differential expression analysis was conducted under default settings with significance thresholds set at (log_2_FC) ≥ 1 and p-value < 0.05 (Tang et al. 2017).

## RESULTS AND DISCUSSION

### ADME and Pharmacokinetic Properties of Stilbene Compounds

The Lipinski and Veber rule-based filter was applied to screen out compound with drug likeliness and better bioavailability profiles. Additionally, ADMET properties such as Absorption, Distribution, Metabolism, Excretion, Toxicity, BBB permeability and GI absorption were also evaluated using Swiss ADME, as shown in Table 1.

**Table 1:** ADME and Pharmacokinetic Properties of Stilbene Compounds using Swiss ADME.

| Molecules | GI absorption | BBB permeant | Lipinski violations | Veber violations | Bioavailability Score | Synthetic Accessibility |
| --- | --- | --- | --- | --- | --- | --- |
| Resveratrol | High | Yes | 0 | 0 | 0.55 | 2.02 |
| Piceid | High | No | 1 | 0 | 0.55 | 4.82 |
| Astringin | Low | No | 1 | 1 | 0.55 | 4.86 |

Regarding GI absorption, Resveratrol demonstrated favourable properties with high gastrointestinal (GI) absorption, blood-brain barrier (BBB) permeability, and no violations of Lipinski’s or Veber’s rules, indicating good oral bioavailability and drug-likeness. It also exhibited a bioavailability score of 0.55 and a low synthetic accessibility score of 2.02, suggesting ease of synthesis. In contrast, Piceid showed high GI absorption but lacked BBB permeability, with one Lipinski’s rule violation and zero Veber violations, indicating acceptable flexibility and polar surface area, and a bioavailability score of 0.55. However, its synthetic accessibility score was higher (4.82), suggesting increased complexity in chemical synthesis. Astringin, on the other hand, exhibited low GI absorption and no BBB permeability, with one violation each of Lipinski’s and Veber’s rules, which may negatively impact its oral bioavailability. Despite this, its bioavailability score remained moderate (0.55). The synthetic accessibility score of 4.86 indicates that it may be synthetically more challenging to produce compared to RVT. Overall, RVT emerged as the most pharmacokinetically favourable candidate among the three compounds, with strong oral absorption potential, CNS penetration, and ease of synthesis.

#### Toxicity Analysis of Resveratrol

The toxicity profile of resveratrol was evaluated using the ProTox-3.0 prediction platform. The compound exhibited a predicted oral LD□ □ of 1560 mg/kg, corresponding to Toxicity Class 4, which is considered “harmful if swallowed” but indicates relatively low acute toxicity. Figure 1 showed, organ-specific toxicity predictions indicated that resveratrol is inactive for hepatotoxicity, neurotoxicity, and respiratory toxicity, but showed a moderate probability of nephrotoxicity and cardiotoxicity. Toxicological endpoint predictions classified resveratrol as inactive for carcinogenicity, immunotoxicity, mutagenicity, cytotoxicity, and clinical toxicity, suggesting a favourable safety profile. Metabolic enzyme interaction analysis revealed that resveratrol is predicted to be an active substrate/inhibitor for CYP1A2, CYP2C9, and CYP3A4, while inactive against CYP2C19, CYP2D6, and CYP2E1. These findings suggest that resveratrol may undergo metabolism through key cytochrome P450 enzymes, particularly CYP3A4, which could influence its bioavailability and potential drug–drug interactions. Overall, these results indicate that resveratrol displays low predicted systemic toxicity, limited organ-specific risks (notably nephrotoxicity and cardiotoxicity), and no major carcinogenic or mutagenic potential. Its predicted interactions with multiple CYP isoenzymes and nuclear receptors further highlight the need for careful pharmacokinetic and drug–drug interaction studies in clinical settings.

**Figure 1:**
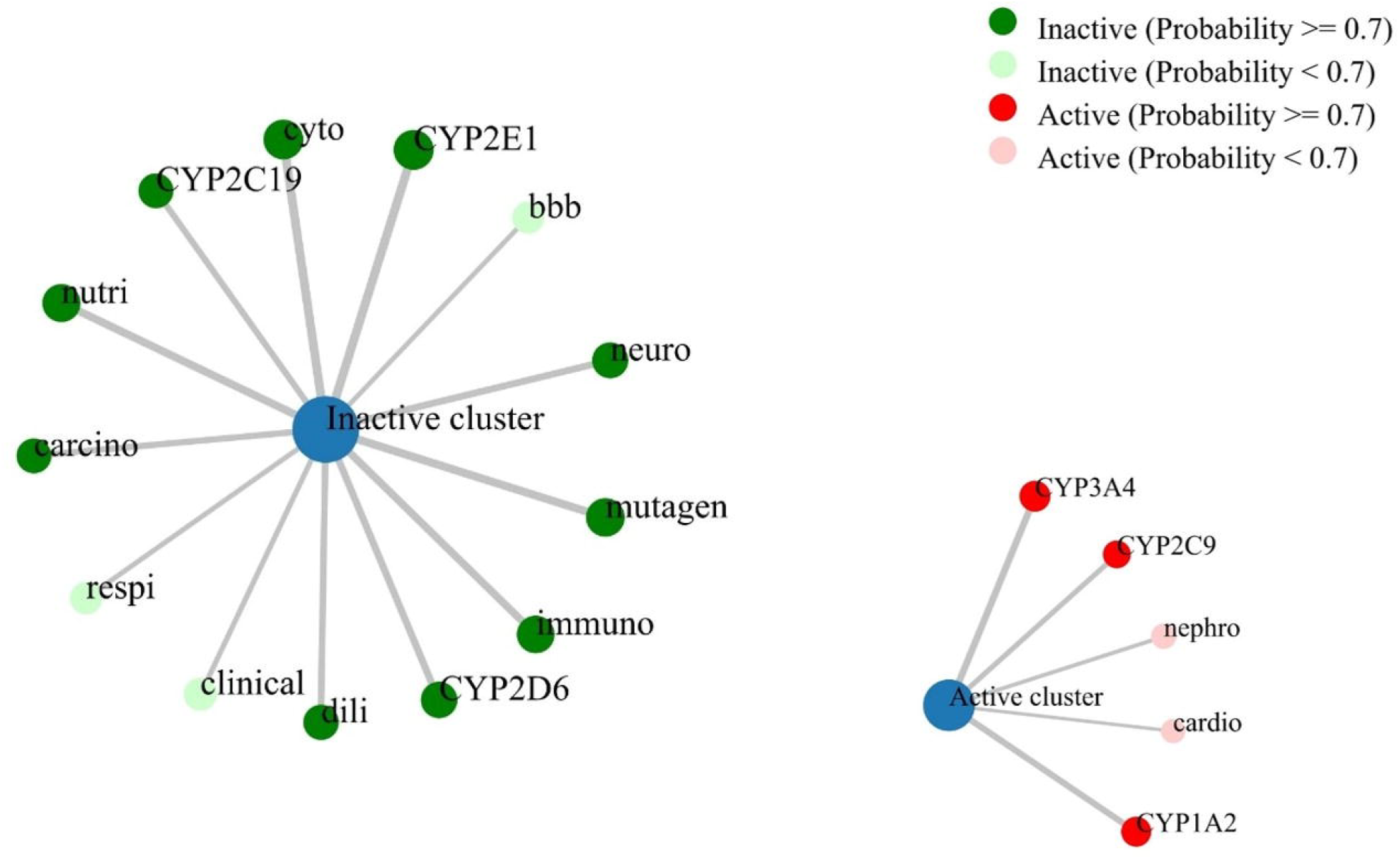
Toxicity Network analysis of Resveratrol using Pro-Tox-III WEB tool. The network chart is intended to the connection between the Resveratrol and Organ Toxicity, Toxicity end points, Metabolism.

### Screening and identification of potential targets of Resveratrol and Pancreatic Cancer

Initially, 100 possible targets for Resveratrol were discovered using the Swiss target prediction database. In all, 1447 known Pancreatic Cancer targets were selected from the Gene Cards and NCBI Gene databases based on a 10 or >10 higher relevance score . Gene symbols having a protein-coding function were chosen to efficiently filter out inconsequential genes from the Gene Cards database. The expected targets of compounds could be supported by the literature evidence. To identify the potential therapeutic targets of resveratrol in pancreatic cancer, we first compiled a dataset of 1,447 known pancreatic cancer-associated targets and 100 predicted resveratrol targets. These two datasets were integrated and analyzed using the Venny tool to visualize their intersections. As shown in Figure 2, a total of 39 genes were found to be shared between resveratrol and pancreatic cancer. These overlapping genes represent the putative intersection where the pharmacological activity of resveratrol may converge with the molecular mechanisms underlying pancreatic cancer pathogenesis. The identification of these 39 common targets is particularly significant since it demonstrates resveratrol’s potential significance in altering crucial pathways involved in pancreatic cancer growth. This overlap provides a narrowed list of possible genes for further inquiry by reducing the large set of disease-associated genes to a group specifically linked to resveratrol. These targets can be used as the foundation for further functional enrichment analysis, network pharmacology research, and mechanistic validation trials, ultimately leading to a better understanding of how resveratrol may exert anti-cancer effects in pancreatic cancer.

**Figure 2:**
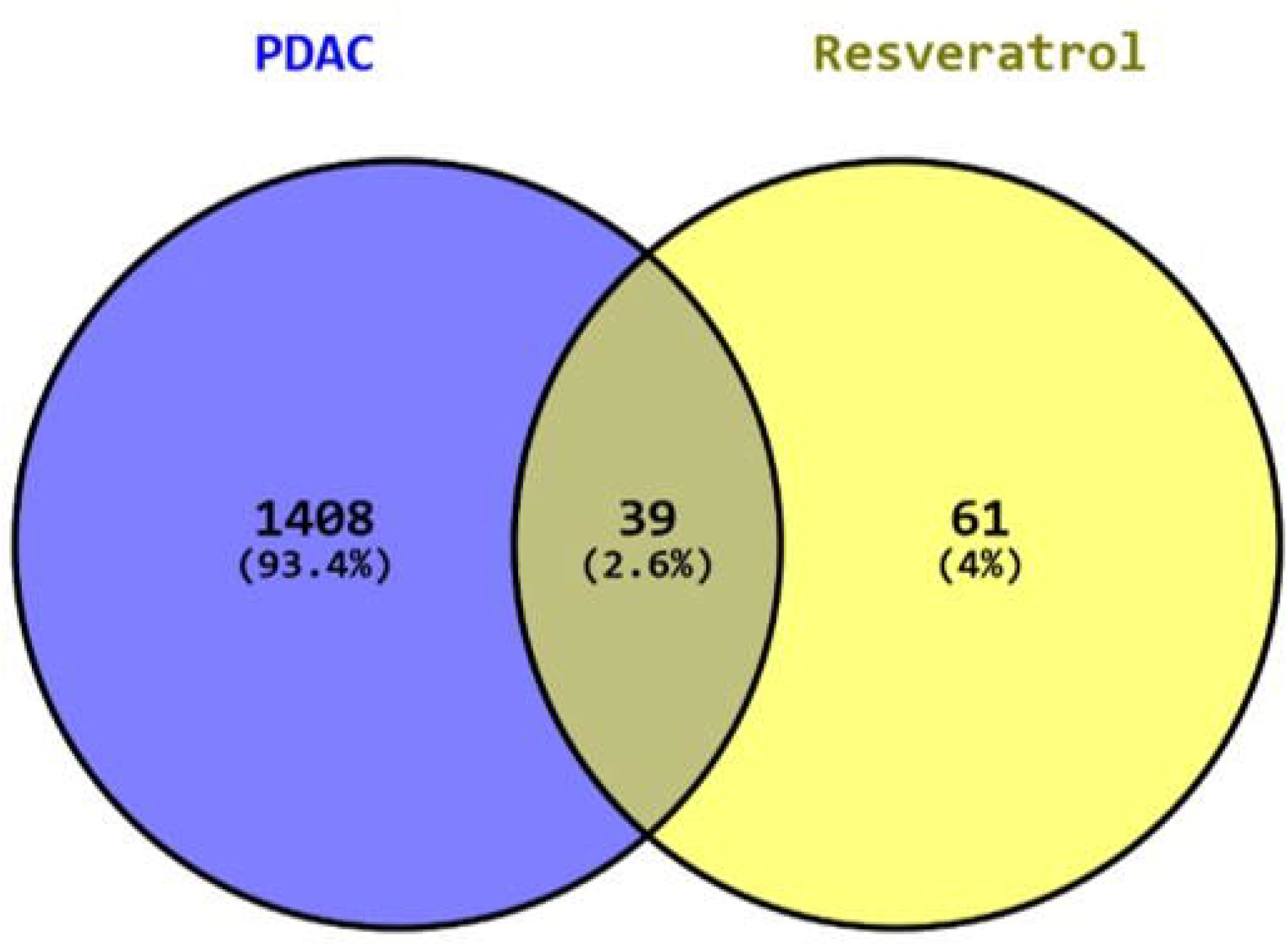
Identification of common targets of Resveratrol and Pancreatic Cancer: Venn diagram illustrating the common targets between pancreatic cancer and resveratrol. The blue circle represents pancreatic cancer-associated targets, while the yellow circle denotes resveratrol-related targets.

### Protein-Protein Interaction (PPI) network construction of drug-potential targets

The 39 common targets identified by overlapping were then used to create a Protein-Protein Interaction (PPI) network of resveratrol-associated targets in pancreatic cancer using the STRING database, as shown in Figure 3a (Supplementary File 1). The confidence score was set to 0.700. Following screening, all targets were found as interacting with other proteins. The average node degree was 9.74, with 190 edges found in the network. Each node’s color corresponds to its degree, with darker nodes representing higher degrees. Edge betweenness correlates with node size, with larger nodes having a higher value. Several proteins, including EGFR, SRC, BCL2L1, PTGS2 PI3KCA, PI3KCB and MTOR, were identified as hub nodes (Figure 3b). These seven hub genes were identified using the CytoHubba plugin in Cytoscape based on betweenness centrality, node degree, and closeness, indicating their critical role in modulating the pharmacological effects of resveratrol (Supplementary file 2). These genes serve as essential regulators in the protein-protein interaction (PPI) network, highlighting prospective targets for resveratrol’s therapeutic benefits. Functional modules were evident within the network: apoptosis and cell survival regulation (BCL2L1, RELA, IGF1R, MTOR), signal transduction and oncogenic pathways (EGFR, RAF1, PI3KCA, KIT, FLT3), drug metabolism and transport (CYP1A2, CYP1B1, ABCC1, ABCB1), and (iv) extracellular matrix remodelling (MMP1, MMP2, MMP9). The high degree of interactions within the PI3K–Akt–mTOR signaling and RAF–MAPK cascade indicates that resveratrol may influence pancreatic tumor progression through the suppression of proliferative and survival pathways. Moreover, the involvement of the NF-κB axis (RELA) highlights the potential anti-inflammatory and pro-apoptotic contributions of resveratrol. Overall, this systems-level analysis suggests that resveratrol exerts its anticancer activity against pancreatic cancer by multi-target modulation of key oncogenic signaling pathways.

**Figure 3a:**
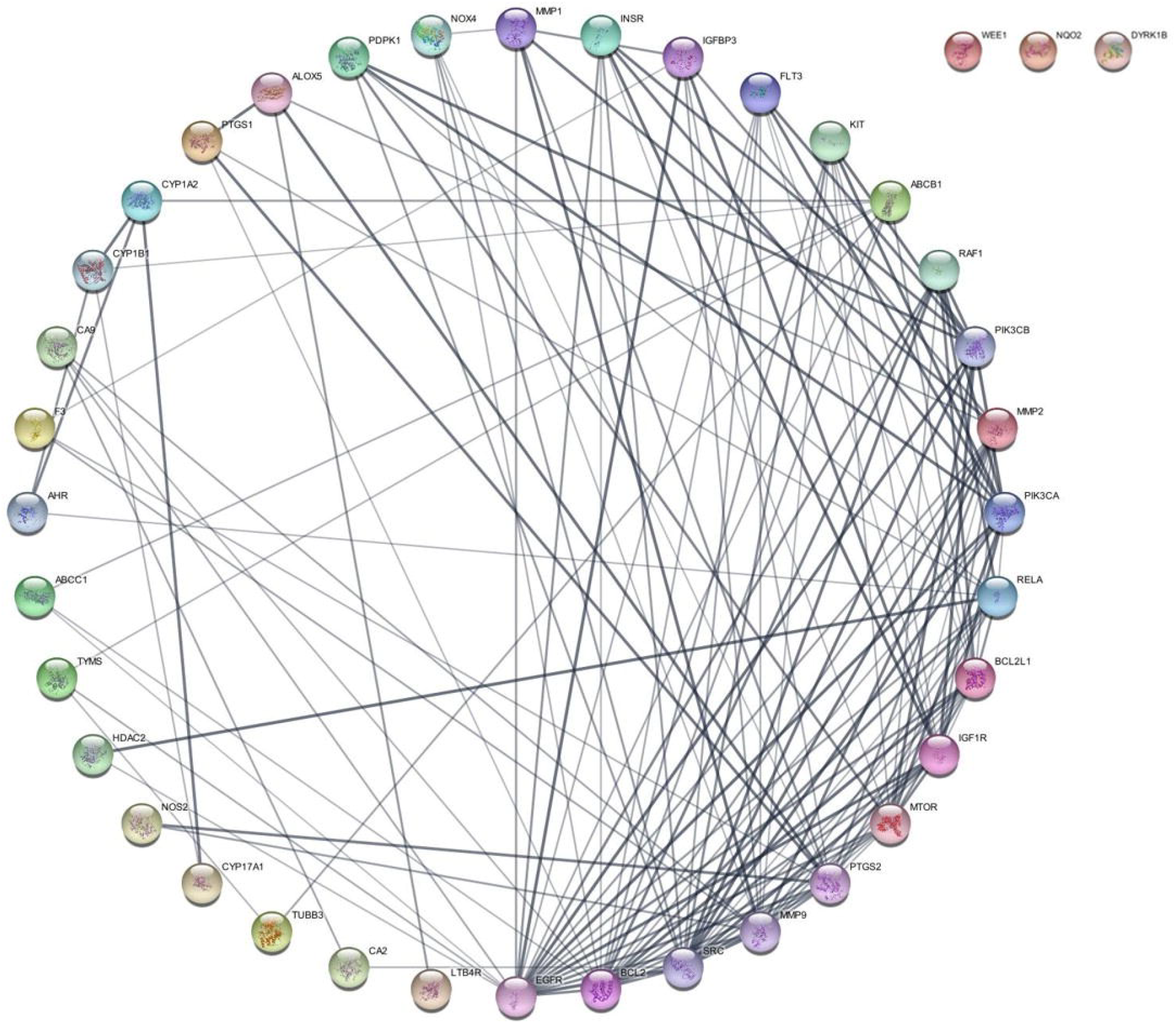
Protein–protein interaction (PPI) network of resveratrol-associated targets in pancreatic cancer. The network comprises 39 nodes and numerous edges. The dense connectivity indicates that resveratrol may exert therapeutic effects through multi-target regulation.

**Figure 3b:**
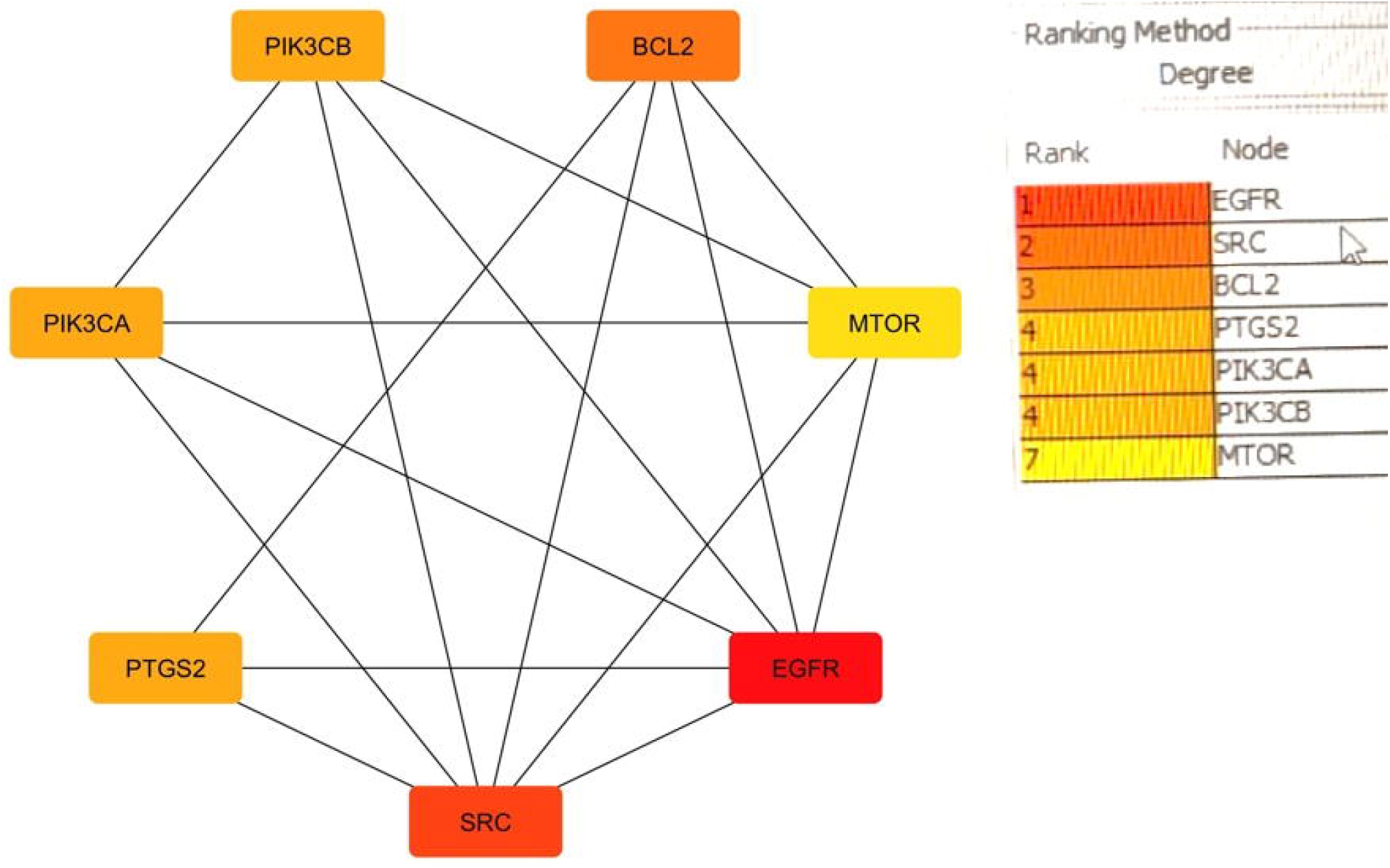
Diagram Representing the hub genes network, seven hub genes such as EGFR, MTOR, PI3KCA, PI3KCB, MTOR, SRC, PTGS2 and BCL2L1 identified using CytoHubba of Resveratrol and Pancreatic cancer.

#### Compound–target interaction network

A compound–target interaction network was constructed in Cytoscape to visualize the relationships between resveratrol and its identified targets, as depicted in Figure 4. A compound-disease target interaction network was created to investigate the link between resveratrol and pancreatic cancer. During the initial screening, 39 overlapping genes were identified as common targets for resveratrol and pancreatic cancer. These genes included significant oncogenes, tumor suppressors, and signaling regulators like EGFR, SRC, BCL2, PTGS1, PTGS2, PIK3CA, PIK3CB, MTOR, RAF1, and RELA, which are known to play important roles in cancer development and survival pathways. Furthermore, several genes associated with angiogenesis and extracellular matrix remodeling growth factor signaling and xenobiotic/drug metabolism were identified, demonstrating resveratrol’s pleiotropic biological activities. The interaction network highlights the ability of resveratrol to target multiple interconnected signaling pathways, including PI3K/Akt/mTOR, EGFR/MAPK, BCL2-mediated apoptosis regulation, and NF-κB axis, thereby providing a systems-level perspective of its potential anticancer activity. The multi-target interaction pattern suggests that RVT may act synergistically by simultaneously modulating tumor growth, angiogenesis, inflammation, drug resistance, and apoptosis regulation in pancreatic cancer.

**Figure 4.**
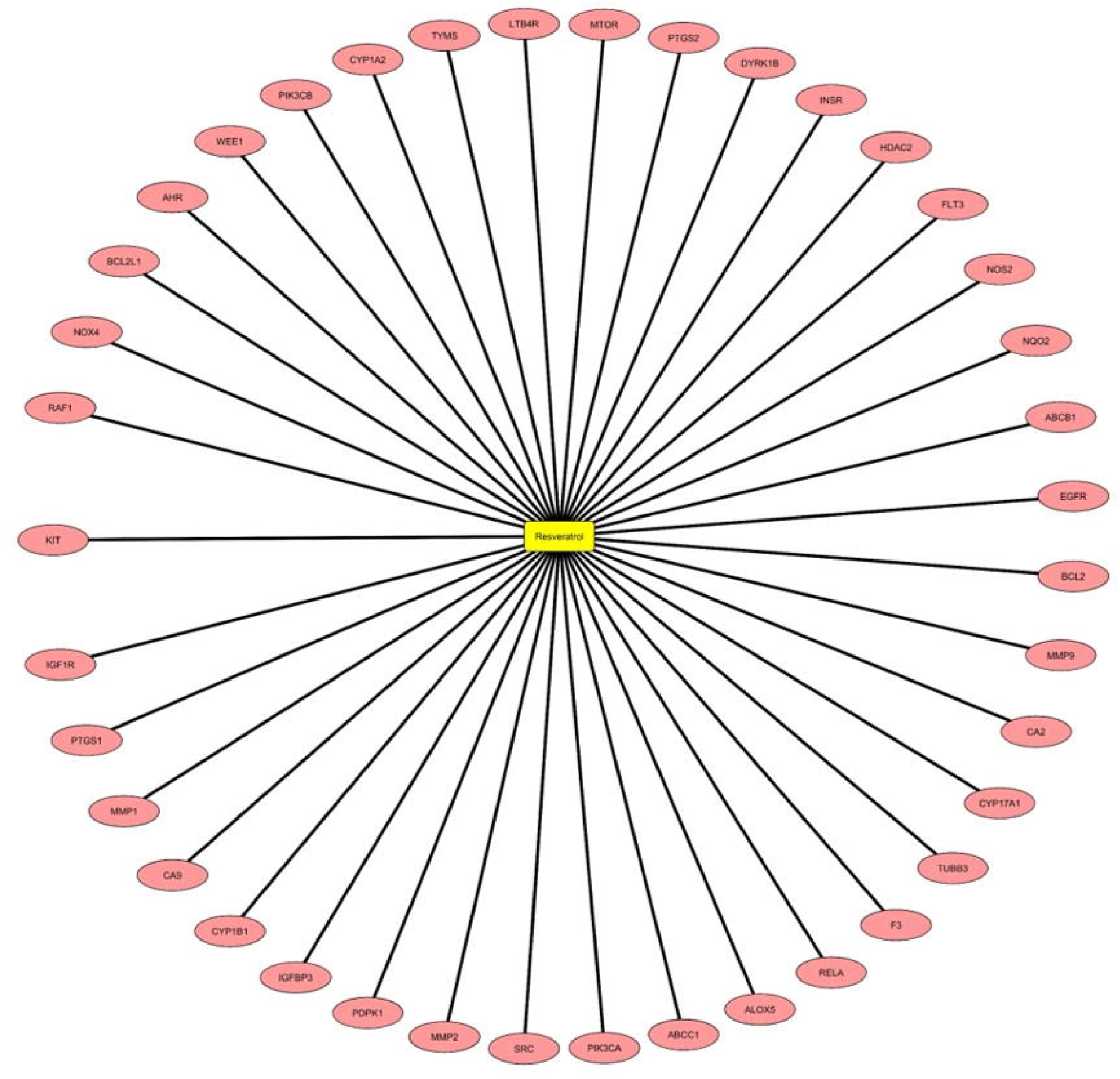
Disease target-compound network. Target-compound network showed the 39 common targets of Resveratrol and Pancreatic Cancer. Target network of resveratrol against pancreatic cancer-associated genes. The network diagram illustrates the multi-target interactions of resveratrol (central yellow node) with several key proteins (pink nodes) implicated in pancreatic cancer progression. The compound demonstrates interactions with critical signaling molecules such as EGFR, MTOR, PI3KCA, SRC, PTGS2 and BCL2, indicating its potential to modulate multiple oncogenic pathways including cell proliferation, apoptosis, and survival signaling cascades.

#### Integration of previously experimentally validated genes

Three reported genes, KRAS, TP53, and CDKN2A, which are associated with pancreatic cancer, were included in addition to computationally identified hub genes (Supplementary Figure 1). KRAS mutations are recognized as one of the earliest genetic events during pancreatic intraepithelial neoplasia (PanIN) progression. The tumor suppressor gene CDKN2A (p16), encoding a cell-cycle regulator, is reported to be inactivated in over 90% of pancreatic cancer cases. Likewise, TP53 mutations, present in approximately 75% of patients, are often characterized by intragenic mutations and loss of heterozygosity, leading to impaired DNA damage responses, cell-cycle arrest, and apoptosis regulation. These genes are together important oncogenic drivers and tumor suppressors in pancreatic cancer [22].

### Analysis of Gene ontology (GO) functional enrichment and potential KEGG target-related pathway

To analyse GO and KEGG pathway functional enrichment, Shiny GO v0.85 and DAVID database was used. The common intersecting genes was entered into the Shiny GO online server and selecting “Human” as the organism. Using a stringent cutoff (FDR < 0.005) This database showed total 560 biological processes, 44 molecular activities, and 20 cellular components for the top 7 hub genes (Figure 5), which displays the top 20 outcomes of the gene enrichment study.

Gene Ontology (GO) enrichment analysis using a stringent cutoff (FDR < 0.005) identified 560 significantly enriched Biological Process (BP) terms. The most enriched pathways included cellular response to decreased oxygen levels (GO:0036294, FDR = 9.98 × 10□ □), negative regulation of apoptotic process (GO:0043066, FDR = 9.98 × 10□ □), negative regulation of programmed cell death (GO:0043069, FDR = 9.98 × 10□ □), and cellular response to hypoxia (GO:0071456, FDR = 9.98 × 10□ □). These enriched biological processes strongly implicate the involvement of the input gene set in cell survival, apoptosis regulation, and hypoxia response pathways [Figure 5a]. These terms predominantly cluster around apoptosis regulation, hypoxia/oxygen response, and cell survival mechanisms. The most highly enriched terms included cellular response to decreased oxygen levels, cellular response to hypoxia, and response to oxidative stress, suggesting that the input genes are tightly associated with oxygen-sensing and stress-adaptive Signaling pathways. Additionally, processes such as negative regulation of apoptotic process, negative regulation of programmed cell death, and anoikis highlight the role of these genes in cell fate determination and resistance to apoptosis, a hallmark of oncogenic transformation and therapeutic resistance. Other enriched processes such as angiogenesis, epithelial cell proliferation, and PI3K-Akt–mediated Signaling regulation reflect the strong involvement of this gene set in tumor progression, survival Signaling, and adaptation to the tumor microenvironment. Collectively, these findings suggest that the resveratrol–target gene network modulates processes central to cell survival, stress adaptation, and oncogenic Signaling in pancreatic cancer.

**Figure 5a:**
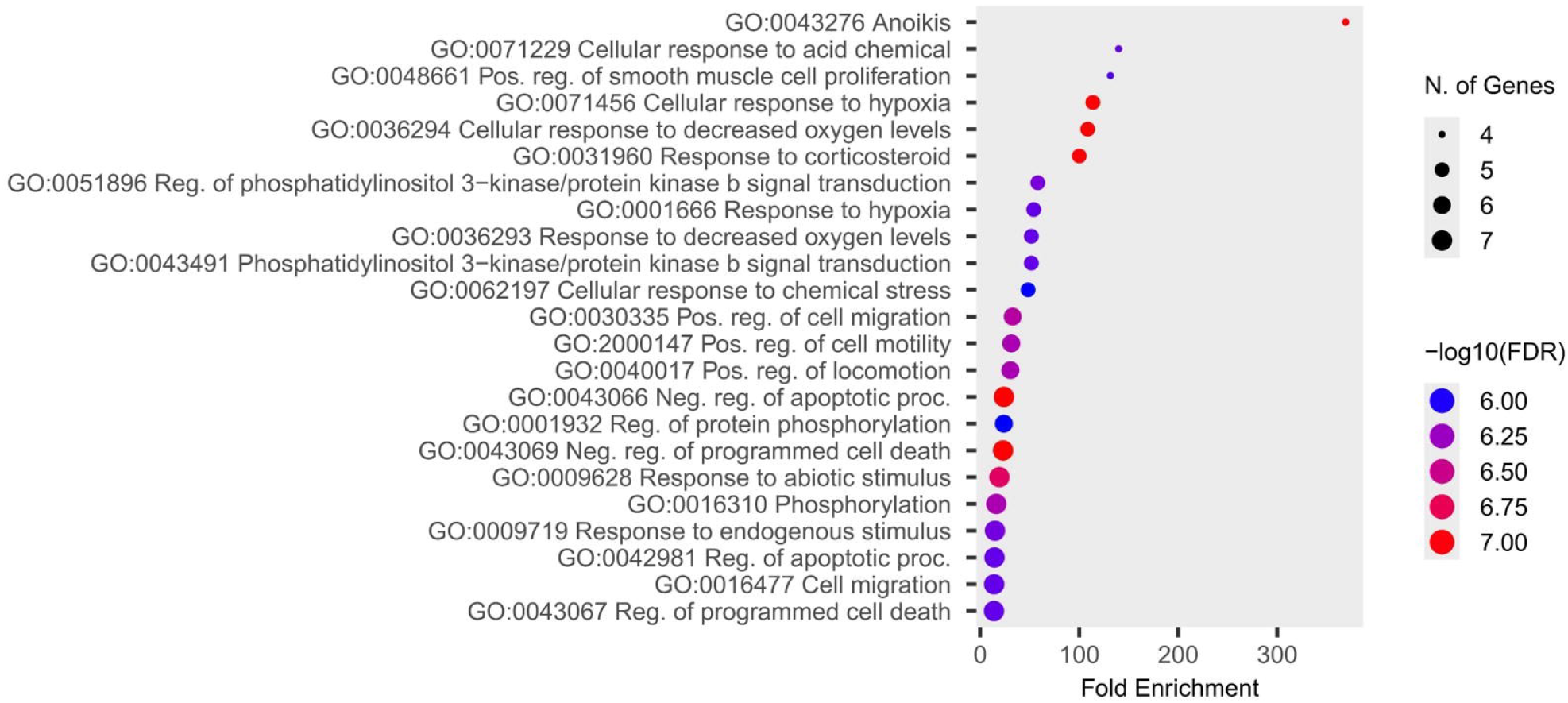
Gene ontology enrichment analysis. GO analysis of the top seven hub genes, EGFR, MTOR, PI3KCA, PI3KCB, MTOR and BCL2L1 using a stringent cutoff (FDR < 0.005) in terms of Biological Process (BP).

Gene ontology (GO) enrichment analysis was performed using ShinyGO on the molecular function (MF) category. Out of the analyzed gene set, 44 unique molecular functions were identified as significantly enriched, with a false discovery rate (FDR) less than 0.005 [Figure 5b]. The top enriched molecular activities included protein kinase activity, kinase activity, phosphotransferase activity with alcohol group as acceptor, various phosphatidylinositol-kinase activities, ATP binding, and nucleotide binding. These functions were represented by key genes such as SRC, MTOR, EGFR, PIK3CB, and PIK3CA, indicating strong involvement in signal transduction and phosphorylation pathways. these findings imply that the resveratrol–target gene network, promotes kinase-related activities and essential signaling nodes to inhibit pancreatic cancer progression, consistent with the enrichment of kinase and phosphorylation-related molecular functions. Additionally, the enrichment of protein tyrosine kinase activity and protein/serine/theonine/ tyrosine kinase activity suggests that resveratrol may interfere with oncogenic signaling cascades mediated by receptor tyrosine kinases such as EGFR.

**Figure 5b:**
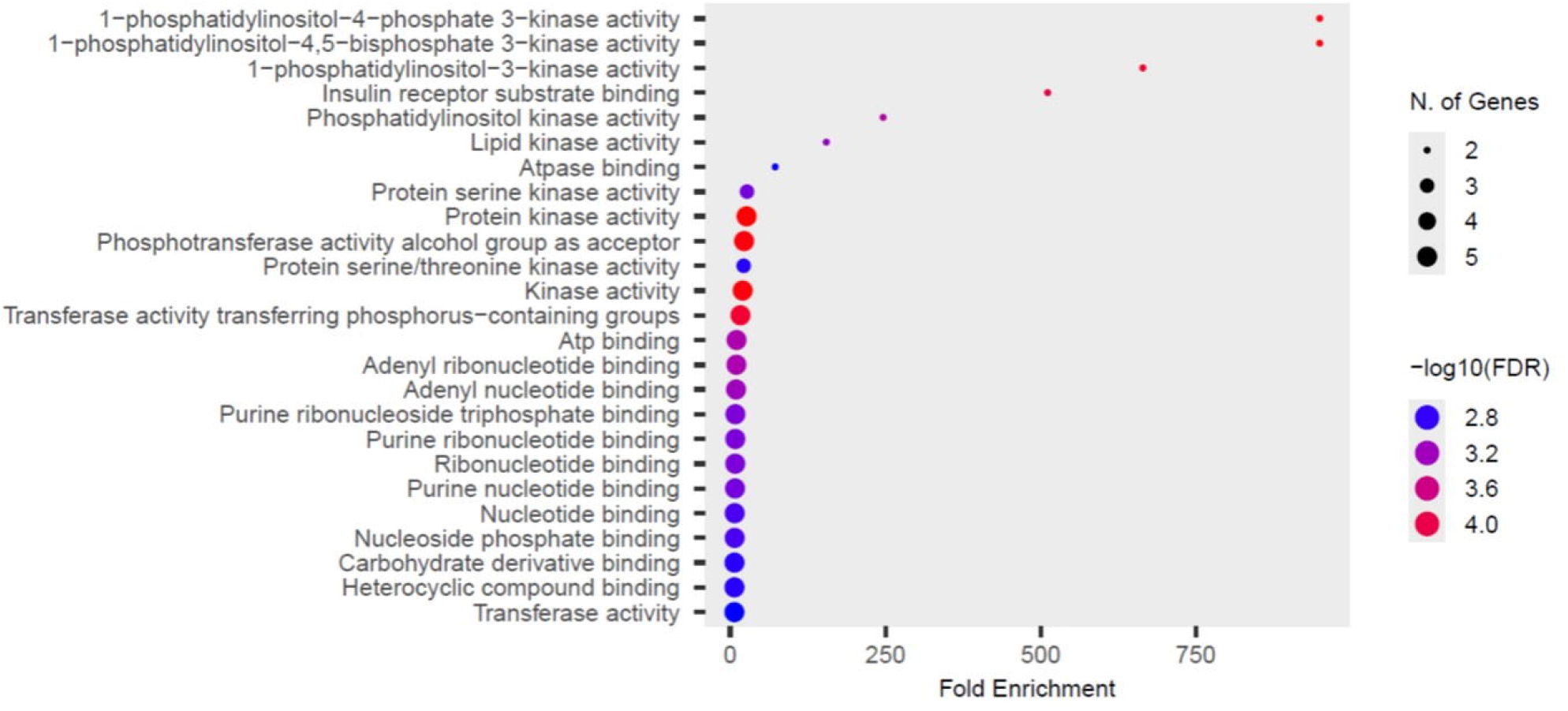
Gene ontology enrichment analysis. GO analysis of the top seven hub genes, EGFR, MTOR, PI3KCA, PI3KCB, MTOR and BCL2L1 using a stringent cutoff (FDR < 0.005) in terms of molecular function (MF).

The cellular component analysis using ShinyGO identified 20 significantly enriched cellular components at an FDR threshold below 0.005. These include critical membrane-associated complexes such as the phosphatidylinositol 3-kinase complex, membrane rafts, microdomains, and organelle envelopes, as well as nuclear components like the nuclear envelope and nuclear membrane [Figure 5c]. Result revealed that all hub genes demonstrating prominent localization patterns relevant to signal transduction, apoptosis, and membrane organization. Key enriched components include the phosphatidylinositol 3-kinase (PI3K) complex and the insulin receptor complex, both of which are integral to signaling pathways that regulate cell growth, metabolism, and survival—processes frequently dysregulated in pancreatic cancer. The presence of genes localized to caveolae, and membrane rafts suggests resveratrol’s involvement in modulating lipid raft–associated signaling, which plays a role in receptor clustering and signal transduction. Additionally, enrichment in the receptor complex supports its potential to disrupt receptor-mediated oncogenic pathways. The association with the organelle outer membrane and nuclear membrane further implies roles in regulating mitochondrial function, nuclear signaling, and apoptosis. These cellular localizations underscore resveratrol’s ability to influence multiple compartments involved in cancer cell signaling and homeostasis.

**Figure 5c:**
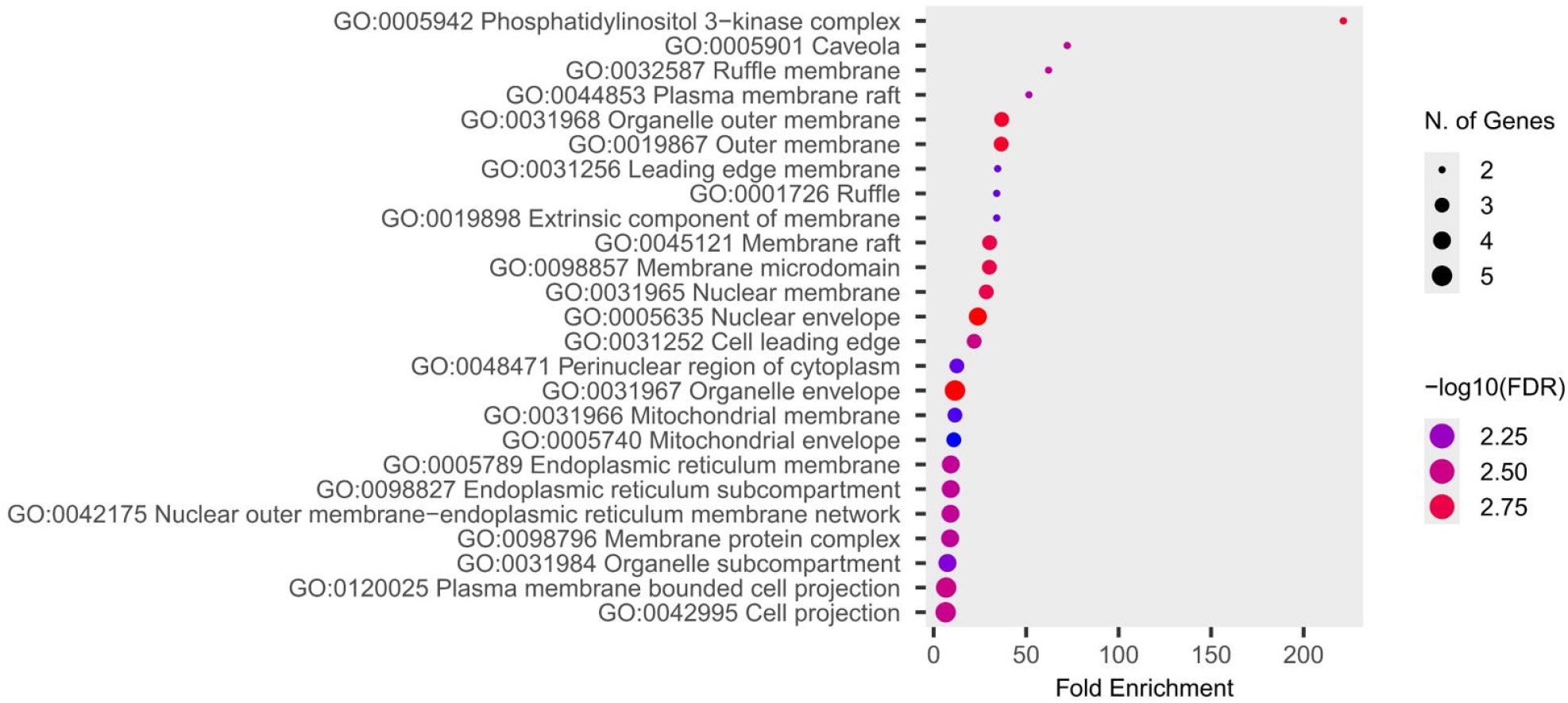
Gene ontology enrichment analysis. GO analysis of the top seven hub genes, EGFR, MTOR, PI3KCA, PI3KCB, MTOR and BCL2L1 using a stringent cutoff (FDR < 0.005) in terms of cellular components (CC).

#### Analysis of the target□pathway network

Proteins and genes work together in interactive and adaptable pathways to carry out pharmacological and biological tasks. KEGG pathway enrichment analysis (FDR < 0.005) identified multiple keys signaling cascades among the overlapping targets, as illustrated in Figure 6a (Supplementary file 3). The database revealed 117 KEGG pathway records with the pancreatic cancer pathway emerging as particularly relevant. The four most significant hub genes, EGFR, MTOR, PIK3CA and PIK3CB, were found to be engaged in the pancreatic cancer pathway out of all seven hub genes. Particularly it has been showed, pancreatic cancer pathway integrates 9 interconnected signaling routes, including the 6 major pathways PI3K–Akt, ErbB, JAK–STAT, MAPK, VEGF, and apoptosis, along with pathways related to, cell cycle regulation, and the p53 signaling cascade. the ErbB Signaling pathway involves 5 hub Genes (EGFR, MTOR PIK3CA, PIK3CB, SRC), The PI3K–Akt Signaling pathway involves 5 hub Genes (EGFR, MTOR, PIK3CA, PIK3CB, BCL2), JAK–STAT Signaling pathway involves 5 hub Genes (EGFR, MTOR, PIK3CA, PIK3CB, BCL2), the VEGF involves 4 hub Genes (PIK3CA, PIK3CB, PTGS2, SRC), the apoptosis pathway involves 3 hub Genes (PIK3CA PIK3CB BCL2) and the MAPK involves 1 hub Genes (EGFR). These hub genes are associated with the pancreatic cancer pathway and play an essential role in its treatment. Moreover, The KEGG phylogenetic association analysis identified clusters of considerably enriched pancreatic cancer pathways based on gene content and functional similarity (Supplementary Fig 2a). Key signaling pathways including PI3K-Akt, ErbB, EGFR tyrosine kinase inhibitor resistance, and JAK-STAT are in close proximity to the pancreatic cancer pathway, highlighting their central roles in tumor cell survival, proliferation, and immune interactions. Furthermore, metabolic pathways, immunological checkpoints, and microenvironment-associated pathways such as focal adhesion and proteoglycans in cancer are all interconnected, demonstrating the comprehensive regulation of pancreatic tumor growth. This clustering supports the complicated network of pathway crosstalk that causes pancreatic cancer. It shows that addressing several interconnected pathways, like with multitarget medicines like resveratrol, can change how tumors behave. Besides, pathways representing other cancer types such as glioma, colorectal cancer, and prostate cancer are also clustered nearby, suggesting shared molecular mechanisms and common oncogenic drivers among these cancers.

**Figure 6a:**
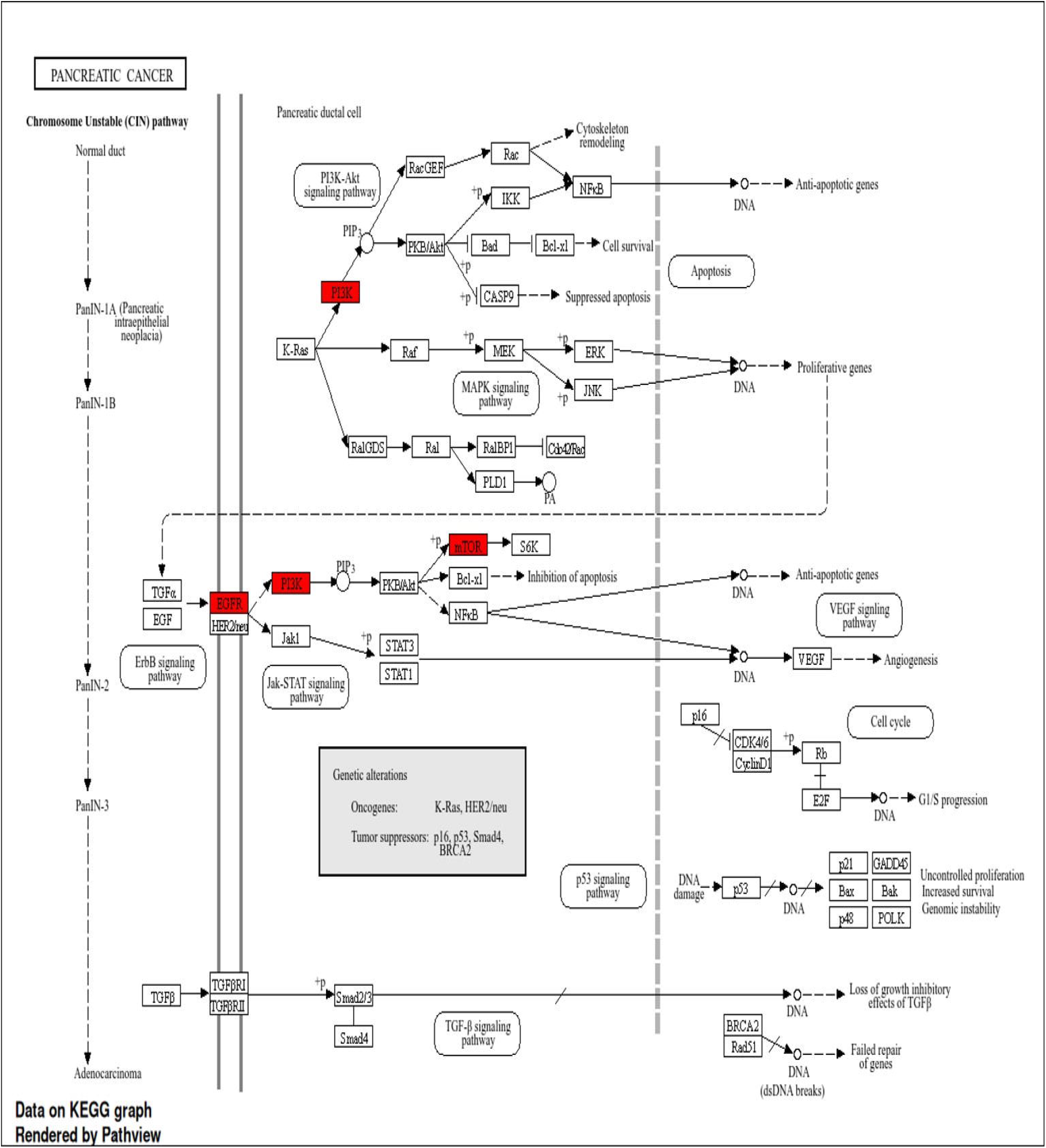
KEGG enrichment network of Resveratrol representing the pancreatic cancer pathway. This pathway contains 5 other pathways, PI3K–Akt-mTOR, ErbB, JAK-STAT VEGF, and MAPK, signaling pathways, that are involved in pancreatic cancer

The KEGG pathway–pathway interaction network analysis, as visualized in the Supplementary Figure 2b highlights the complex interplay among signaling cascades enriched from the overlapping gene targets of resveratrol. Notably, several key pathways—such as the PI3K-Akt signaling pathway, ErbB, EGFR tyrosine kinase inhibitor resistance, HIF-1 signaling pathway, and pathways in cancer—emerge as highly interconnected hubs. These central nodes suggest that resveratrol exerts its effects not through isolated mechanisms but via modulation of a broad network of oncogenic and regulatory processes. The prominence of the “pathways in cancer” node underscores its integrative role, connecting with multiple cancer-specific and signaling-related pathways, including prostate cancer, pancreatic cancer, and microRNAs in cancer.

Additionally, the clustering of pathways related to chemical carcinogenesis, receptor activation, and oxidative stress indicates a potential involvement of resveratrol in influencing inflammatory responses and metabolic reprogramming. Figure 6b illustrates network map which indicates the multi-targeted therapeutic potential of resveratrol, particularly through its regulation of core pathways involved in cancer proliferation, survival, angiogenesis, and resistance mechanisms.

**Figure 6b:**
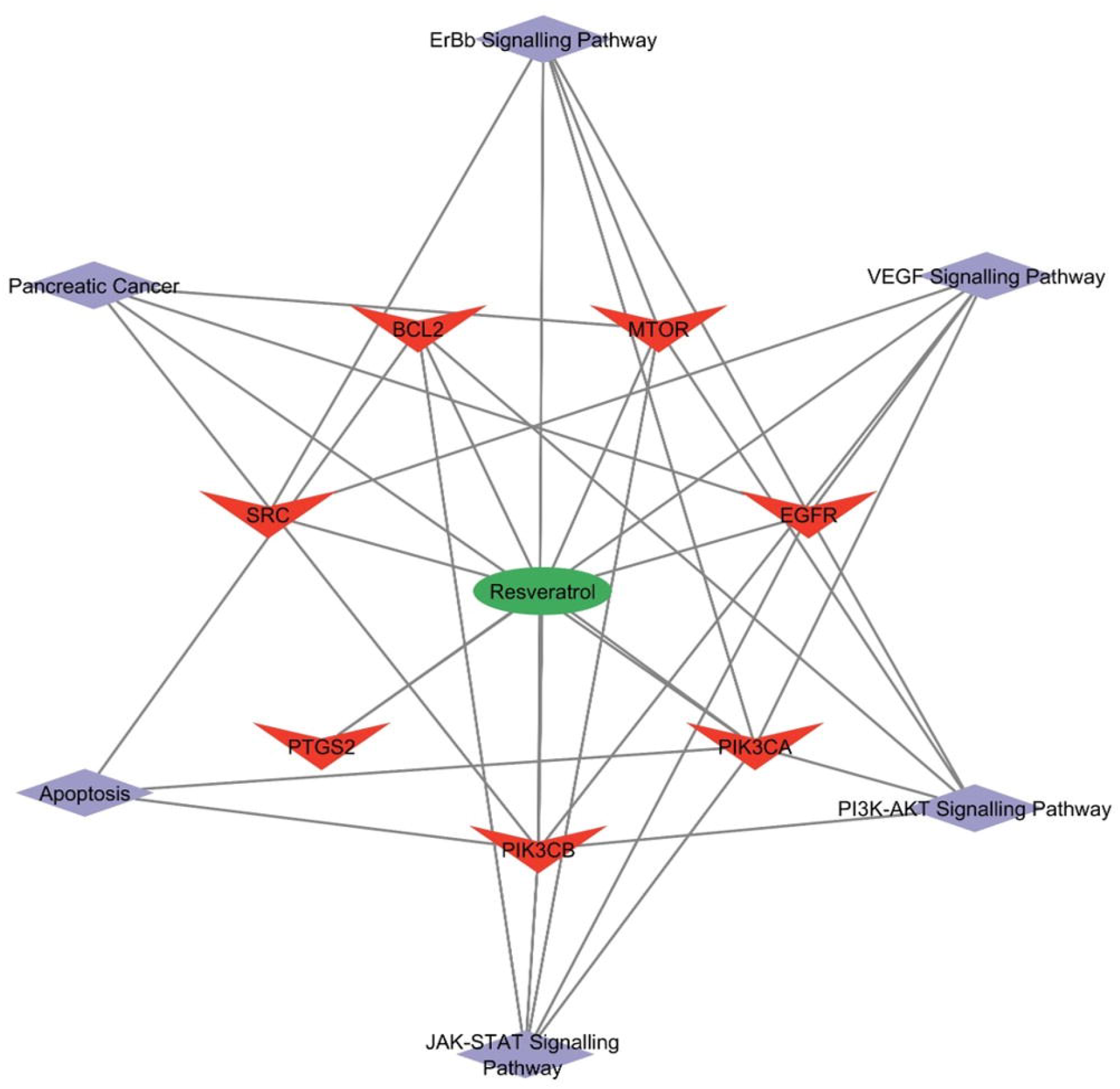
Disease pathway–target–compound network. Disease pathway–target–compound network illustrating the multi-target efficacy of resveratrol against pancreatic cancer. The network diagram visualizes resveratrol (green ellipse) centrally connected to multiple molecular targets (red v-shaped nodes), including BCL2, MTOR, SRC, EGFR, PTGS2, PIK3CA, and PIK3CB. These targets, in turn, are linked to critical disease signaling pathways (purple hexagonal nodes) such as PI3K-AKT, ErbB, JAK-STAT, VEGF, apoptosis, and pancreatic cancer itself. Black lines denote the interaction between resveratrol and respective molecular targets, and between targets and disease pathways, highlighting the compound’s multi-faceted mechanism of action through simultaneous modulation of oncogenic and apoptotic signaling in pancreatic cancer.

#### Molecular Docking Validation

To validate the binding affinity between resveratrol and key protein targets, molecular docking was conducted using AutoDock Vina. Table 2 summarizes the docking results of Resveratrol against ten key protein targets associated with pancreatic cancer. The binding affinity (AutoDock Vina score in kcal/mol), number of hydrogen bonds, and interacting residues are listed for each protein. Figure 7a, 7b and 7c shows the 2D and 3D images of the molecular interaction diagram depicting the interactions of resveratrol with various target proteins. It shows the docking results demonstrated lowest binding energies, with resveratrol showing strong interaction with KRAS (binding affinity = –8.2 kcal/mol) and forming 2 hydrogen bonds and 11 hydrophobic contacts at the active site (Figure 7c). While the other two proteins, i.e., EGFR and MTOR, exhibit moderate binding affinities i.e. -7.9 and -7.7 kcal/mol respectively, and forming 3 hydrogen bonds showing more stability(Figure 7a). Therefore, the binding of all the three proteins with the resveratrol might yield better results in drug design and therapeutic applications because of its strong binding ability.

**Table 2:** Molecular Docking Analysis of Resveratrol with Target Proteins Involved in Pancreatic Cancer.

| S.No. | Protein | Molecule | Vina score | No. of hydrogen bond | Interacting residues |
| --- | --- | --- | --- | --- | --- |
| 1 | BCL2 | Resveratrol | -6 | 0 | VAL B: 153, MET B:112, PHE B:109, PHE B:101, ALA B:146, ASP B:108, GLU B:111, ARG B:107, LEU B:134, PHE B:150, GLU B:149 |
| 2 | EGFR |  | -7.9 | 3 | THR A:790, LEU A:718, LEU A: 844, ALA A: 743, VAL A: 726, GLY A: 796, LEU A:792, MET A: 793, **LYS A:745, THR A :854**, PHE A: 856, GLU A:762, ASP A: 855, **LEU A: 788**, MET A: 766 |
| 3 | MTOR |  | -7.7 | 3 | **GLU B: 2196**, PRO B:01940, ILE B:1943, THR B: 2232, **SER B: 1982**, THR B: 1983, THR B: 1984, LEU B:2426, ALA B: 1979, ARG B: 1987, ARG B:2193, VAL B:1978, TYR B: 1974, ILE B: 2228, LEU B: 2230, **PRO B: 2229** |
| 4 | PIK3CA |  | -7.6 | 2 | **GLU A:81, ARG A:79**, GLU A:80, MET A:123, ARG A:93, GLU A:76, GLN A:75, THR A:74, PRO A:124, GLU A:78, ARG A:115, VAL A:125, ASN A:114, ILE A: 112 |
| 5 | PIK3CB |  | -7.4 | 0 | PHE A:928, GLU A:846, LEU A: 804, ASP A:807, LYS A:799, ASP A:802, ASP A:931, ILE A:845, ILE A:767, VAL A:847, TYR: 833, MET A:920, ILE A: 930, MET A:773, TRP A: 781, VAL A:848 |
| 6 | PTGS2 |  | -7.6 | 0 | LEU A:531, SER A: 353, ARG A: 513, HIS A: 90, ALA A:516, GLN A:192, VAL A: 523, ILE A: 517, PHE A: 518, LEU A: 352, ALA A: 527, VAL A: 349, TRP A: 387, TYR A: 385, TRR A: 348, SER A: 530 |
| 7 | SRC |  | -7.7 | 0 | ALA A: 390, ASP A: 404, MET A: 314, LEU A: 325, THR A:338, VAL A: 323, ILE A: 336, PHE A: 405, ALA A: 403, LYS A: 295, ALA A: 293, TYR A: 340, VAL A: 281, LEU A: 273 ASN A: 391, LEU A: 393 |
| 8 | KRAS |  | -8.2 | 2 | LEU B: 120, **SER B: 145**, ASP B: 119, **LYS** |
|  |  |  |  |  | **B: 147**, CYS B: 118, LYS B: 117, ASN B: 116, PHE B: 28, ALA B: 146, GLY B: 15, ALA B: 18, PRO B: 34, SER B: 17, GLU B:31, TYR B: 32, VAL B: 29, ASP B:30 |
| 9 | TP53 |  | -6.6 | 0 | PRO B: 153, PRO B: 152, GLY B: 154, THR B: 155, VAL B: 147, TRP B: 146, LEU B: 145, ASP B: 228, CYS B: 229, PRO B: 223, THR B:230, CYS B: 220, GLU B: 221, PRO B: 151, THR B: 150, PRO B: 222 |
| 10 | CDKN2A |  | -7 | 1 | ARG A: 168, TYR A: 170, ARG A: 220, LYS A: 216, GLN A: 173, PRO A: 217, TRP A: 184, PRO A: 148, **GLU A: 211**, TYR A: 185, THR A: 106, GLN A: 149, THR A: 182 |

**Figure 7a.**
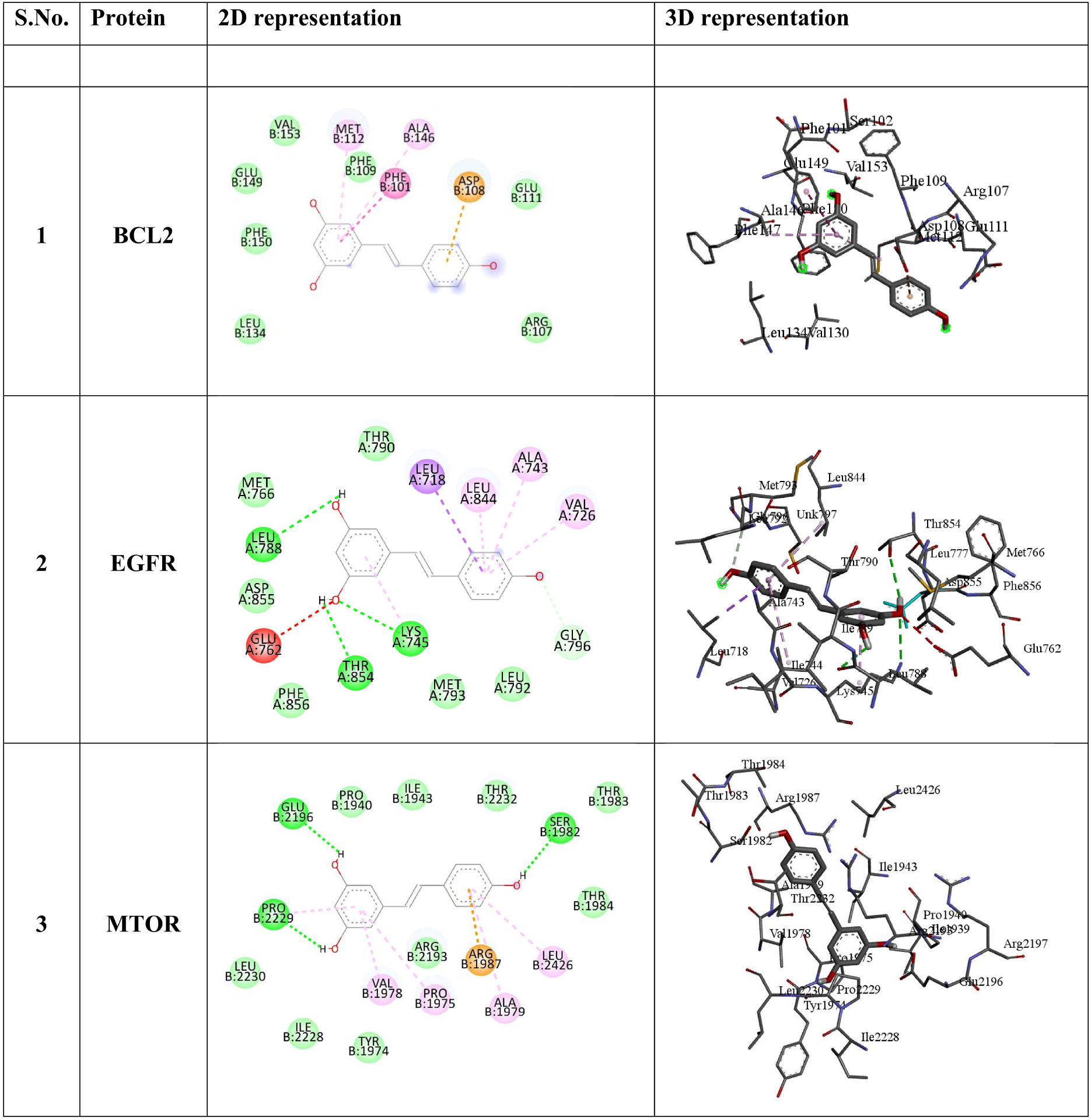
2D and 3D images of the molecular interaction diagram depicting the interactions of resveratrol with BCL2, EGFR and MTOR via AutoDock Vina.

**Figure 7b.**
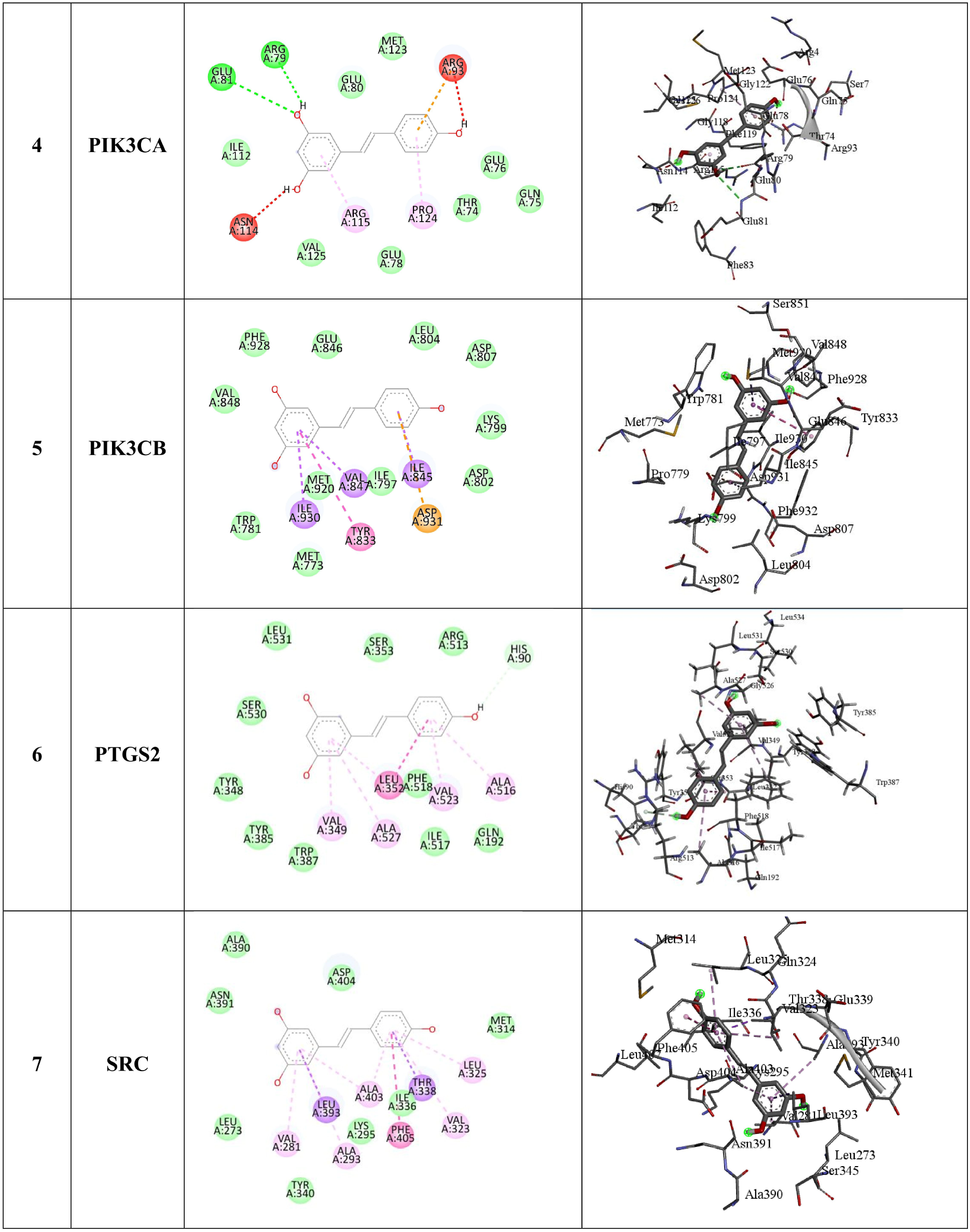
2D and 3D images of the molecular interaction diagram depicting the interactions of resveratrol with PIK3CA, PIK3CB, PTGS2 and SRC via AutoDock Vina.

**Figure 7c.**
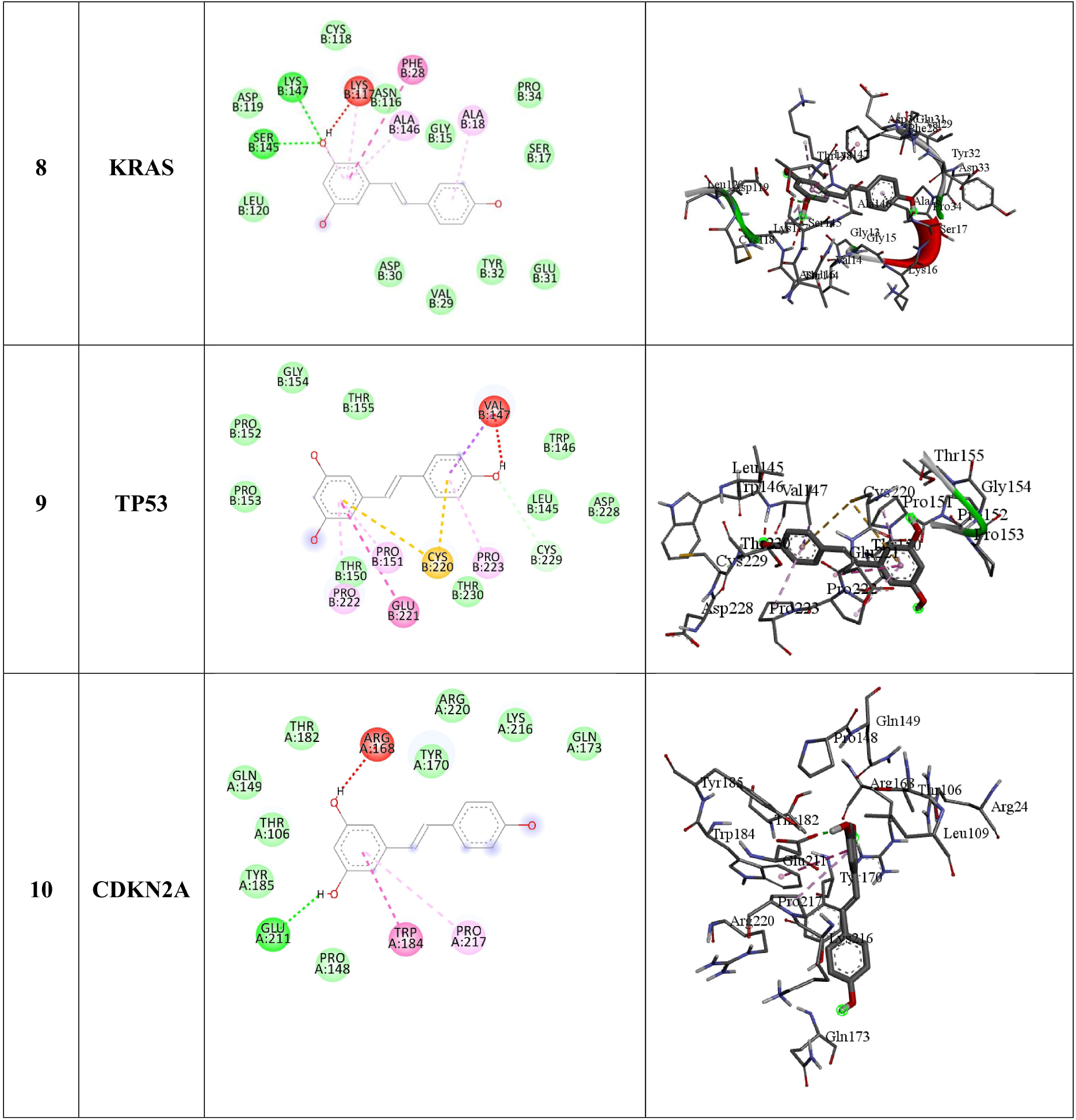
2D and 3D images of the molecular interaction diagram depicting the interactions of resveratrol with KRAS, TP53, and CDKN2A via AutoDock Vina.

### Validation of screened hub genes using mRNA expression

For external validation of the effect of the expression of the screened hub genes on pancreatic cancer, GEPIA2 were employed to assess the mRNA expression. The differential expression of mRNA of the genes, EGFR, BCL2, PIK3CA, PIK3CB, PTGS2 and MTOR were seen to be upregulated in the cancerous tissue (Figure 8). The results of the mRNA expression showed that the hub genes, PIK3CA, PIK3CB, and PTGS2 exhibited markedly higher expression levels in tumor samples, indicating strong upregulation. Similarly, EGFR, BCL2, and MTOR were consistently upregulated, further confirming its upregulation in PAAD. Therefore, these findings highlight the aberrant activation of key oncogenic signaling pathways, underscoring their potential roles in pancreatic cancer and their value as candidate biomarkers or therapeutic targets.

**Figure 8.**
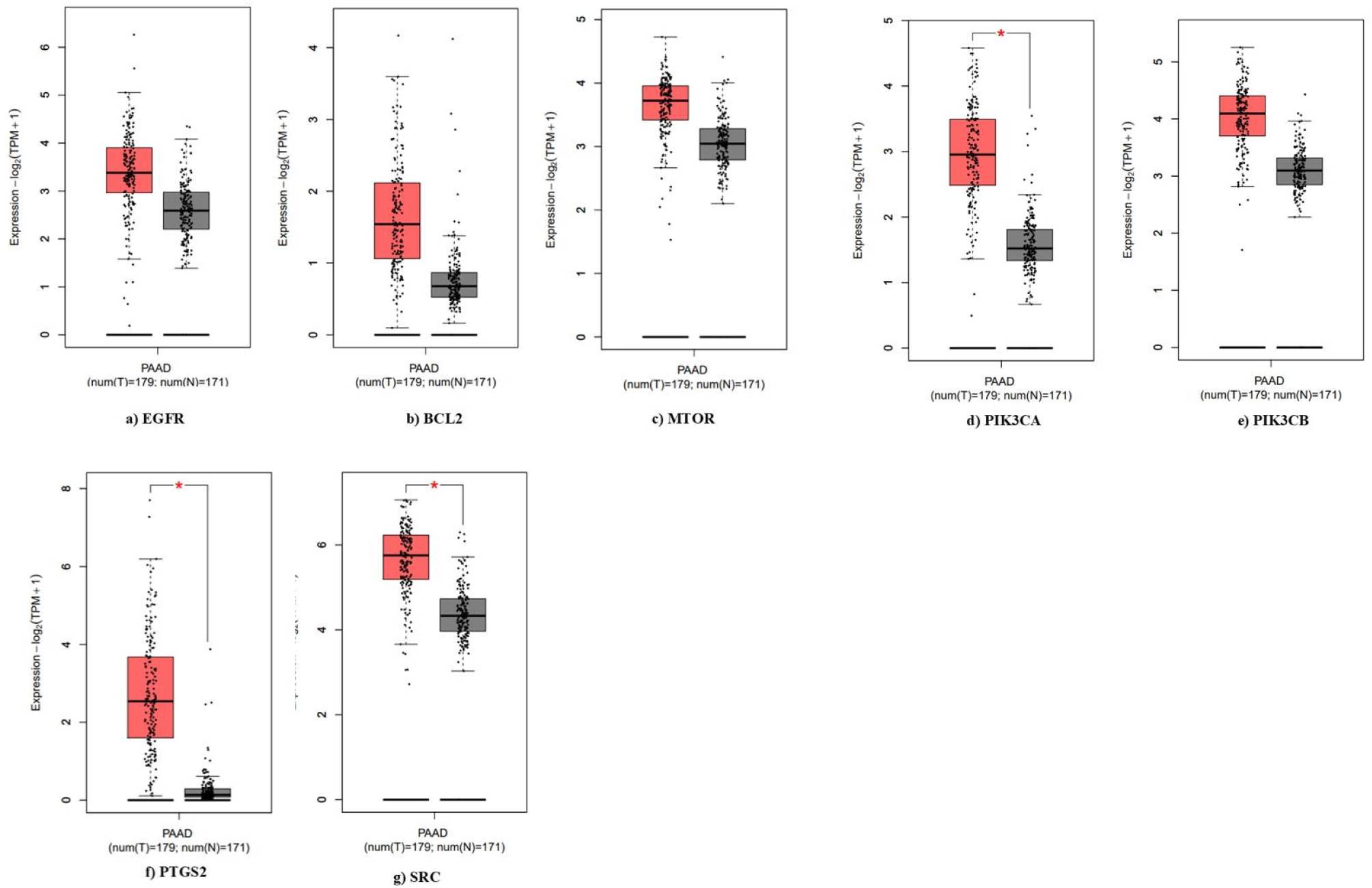
Box plot analysis of Hub gene expression in pancreatic adenocarcinoma (PAAD) tissues versus normal controls using GEPIA 2 software. A significant upregulation of EGFR, BCL2, SRC, PIK3CA, PIK3CB, PTGS2 and MTOR is observed in PAAD tissues compared to normal controls, as indicated by the red asterisk.

## Discussion

Pancreatic ductal adenocarcinoma, the most prevalent form of pancreatic cancer, is associated with various risk factors including smoking, obesity, diabetes, chronic pancreatitis, and reduced physical activity (Zottl et al. 2025). Surgical resection remains the primary curative approach for pancreatic cancer; however, only about 15– 20% of patients are eligible for pancreatectomy due to late-stage diagnosis. Even among those who undergo complete tumor removal, the overall prognosis remains poor (Wong et al. 2023). Besides surgery, synthetic oral chemotherapy drugs are commonly used in clinical management. However, these drugs are often associated with significant adverse effects (Costa et al. 2015. Min and Lee 2022). In recent years, plant-derived natural compounds and their analogs—such as carotenoid (crocetinic acid) (Rangarajan et al. 2015), saponin (sapogenin, diosgenin) (Guo et al. 2019), sesquiterpene (elemene) (Long et al. 2019), apigeniflavan (Das et al. 2023) and kaempferol and catechin (Agrawal et al. 2023) have attracted increasing attention for their potential inhibitory effects against pancreatic cancer progression. Resveratrol (M.W. 228.24), a polyphenolic, stilbenoid compound found in many plants, has received a lot of attention in recent decades for its wide range of therapeutic effects, which include antiangiogenic, immunomodulatory, antimicrobial, neuroprotective, anticancer, antidiabetic, and cardioprotective properties (Zhang et al. 2021). In this comprehensive study, we investigated pharmacokinetic and toxicity, network pharmacology, and molecular docking properties of resveratrol with a specific focus on its therapeutic potential against pancreatic cancer. Initially, pharmacokinetic and toxicity assessments were conducted on the various stilbenoid phytocompounds including resveratrol, piceid, and astringin; resulting in resveratrol being identified as the most pharmacokinetically advantageous option. The ADMET and drug-likeness profiling revealed that resveratrol possesses high gastrointestinal (GI) absorption and blood-brain barrier (BBB) permeability, with no violations of Lipinski’s or Veber’s rules, indicating favourable oral bioavailability and drug-likeness (Daina et al. 2017, Salla et al. 2024). Its low synthetic accessibility score (2.02) also suggests ease of chemical synthesis compared to other compounds i.e. Piceid and Astringin. Resveratrol is known to be rapidly absorbed in both humans and animals, and although it undergoes extensive metabolism, primarily via glucuronidation and sulfation in the liver and intestines, resulting in low systemic availability of the parent compound (Zhang et al. 2021). This reproducible and well-characterized pharmacokinetic footprint provides a strong foundation for its optimization through formulation strategies, thereby supporting its potential as a drug candidate (Ghafouri-Fard et al. 2022, Khattar et al. 2022). In contrast, Piceid, the glucosylated derivative of resveratrol, exhibits slower intestinal absorption, reduced cellular uptake, and a delayed time to reach peak plasma concentration, which diminishes its pharmacological efficiency despite improved solubility (Jiang et al. 2009). Astringin, a glycosylated derivative of piceatannol, has even less pharmacokinetic data available, with limited understanding of its absorption, metabolism, and systemic distribution, making it the least predictable among the three (Jiang et al. 2009). Toxicity profiling using ProTox-III showed that resveratrol demonstrates a low toxicity profile across major parameters, including hepatotoxicity, carcinogenicity, and cytotoxicity, further supporting its safety for potential therapeutic use. In support of our findings, previous studies have consistently demonstrated that resveratrol possesses a favourable safety profile across both preclinical and clinical settings. Even at maximum tolerated doses, oral administration of resveratrol showed no evidence of carcinogenicity, skin or eye irritation, allergenicity, or significant estrogenic activity in vivo (Khattar et al. 2022). Furthermore, high-dose exposure did not adversely affect reproductive performance or bone health, reinforcing its overall tolerability. These observations collectively highlight the broad safety margin of resveratrol and align with its growing consideration as a viable candidate for therapeutic and preventive applications (Zhang et al. 2021). In addition, epidemiological studies indicate a correlation between higher resveratrol intake and reduced risk of breast cancer, while its capacity to cross the blood–brain barrier expands its therapeutic applicability beyond oncology to neurological disorders. Collectively, these findings underscore the low toxicity and broad safety margin of resveratrol strengthening its potential as a promising chemopreventive and therapeutic agent (Khattar et al. 2022). Network pharmacology analysis identified a total of 39 overlapping targets between resveratrol and pancreatic cancer, which were subsequently used to construct a protein-protein interaction (PPI) network. Hub genes such as EGFR, SRC, BCL2, PTGS2, PIK3CA, PIK3CB, and MTOR were identified by network pharmacology analysis as important mediators in the relationship between resveratrol and pancreatic cancer. These genes are known to contribute to chemoresistance, tumor growth, and a poor prognosis in pancreatic cancers (Agrawal et al. 2023). Pancreatic cancer survival is characterized by resistance to apoptosis, which is conferred by overexpression of BCL2, but dysregulated EGFR and SRC signaling encourage unchecked growth and metastasis (Matsuoka 2016). The inflammatory tumor microenvironment, which is closely linked to the advancement of pancreatic cancer. The discovery of PIK3CA, PIK3CB, and MTOR is significant because it highlights the importance of the PI3K/Akt/mTOR pathway, which is often overactive in pancreatic cancers and a major target of resveratrol ‘s modulatory effects (Guo et al. 2019). It has demonstrated that resveratrol may exhibit synergistic anticancer properties by concurrently regulating these hub genes, potentially involving the suppression of angiogenesis and metastasis, growth inhibition, and the activation of apoptosis (Jiang et al. 2009, Zhang et al. 2021).

Gene ontology (GO) and KEGG pathway enrichment analyses indicated that resveratrol targets several biological processes and molecular functions, including oxidative stress response, kinase signaling, receptor tyrosine kinase activity, and metabolic regulation. Angiogenesis, epithelial cell proliferation, and PI3K-Akt-mediated signaling all emphasize the relevance of enriched genes in tumor growth and therapeutic resistance (Jiang et al. 2009). Key pathways including PI3K-Akt, ErbB, JAK-STAT, MAPK, VEGF, and apoptosis were shown to be considerably enriched (FDR < 0.005). All of these play key roles in pancreatic cancer growth, survival, and medication resistance. Since the pathway analysis indicated a significant enrichment of the PI3K-Akt-mTOR axis and other survival pathways in pancreatic cancer, resveratrol’s ability to inhibit these molecular targets gives a compelling mechanistic rationale for its therapeutic potential. By attenuating aggressive features of pancreatic cancer, resveratrol interferes with highly enriched oncogenic pathways, offering a multi-targeted strategy that encompasses both direct modulation of signaling cascades and indirect effects on the tumor microenvironment. Moreover, its capacity to reverse resistance mechanisms highlighted in the KEGG pathway enrichment suggests a promising role in improving therapeutic outcomes (Bertelli 2005, Visioli et al. 2011, Matsuoka 2016). Molecular docking analysis provided supportive evidence for the multi-targeted potential of resveratrol in pancreatic cancer by confirming strong binding affinities with key hub proteins identified through network pharmacology (Agrawal et al. 2023). Among the hub genes, EGFR (–7.9 kcal/mol) and MTOR (–7.7 kcal/mol), both central regulators of survival, growth, and metabolic adaptation in pancreatic tumors, exhibited strong interactions with resveratrol, forming multiple hydrogen bonds that indicate enhanced binding stability and a potential for functional inhibition of these signaling pathways (Ouissam et al. 2024). Resveratrol also demonstrated favourable binding with PIK3CA and TP53, suggesting its capacity to modulate both the PI3K– Akt–mTOR axis and tumor-suppressor networks (Ghafouri-Fard et al. 2022). In addition to the hub genes identified through network pharmacology, key oncogenes and tumor suppressors such as KRAS, TP53, and CDKN2a were considered for molecular docking analysis for comparative observations. These genes are key drivers of pancreatic cancer biology, with KRAS mutations causing aberrant signaling that fuels uncontrolled cell growth and TP53 loss impairing genomic stability and apoptotic responses. Dysregulation of cyclin-dependent kinases (CDKs) further impairs cell cycle control, contributing to tumor development and treatment resistance (Matsuoka, 2016). Interestingly, KRAS, although not identified as a direct target of resveratrol in the enrichment analysis, showed the highest binding affinity (–8.2 kcal/mol). This finding is significant, as mutant KRAS acts as a key oncogenic driver in pancreatic cancer by sustaining aberrant Signaling cascades that promote uncontrolled proliferation and therapeutic resistance (Ghafouri-Fard et al. 2022, Agrawal et al. 2023, Pergolizzi and Brower 2024). Resveratrol seems to change these molecular factors via changing the downstream signaling of mutant KRAS, bringing back apoptotic pathways linked to TP53, or lowering CDK activity to stop the cell cycle. Integrating wet-lab validation with network pharmacology predictions enhances RVT’s mechanistic comprehension, revealing its capacity to target both empirically validated and computationally identified pathways (Agrawal et al. 2023). Several studies have shown that resveratrol downregulates KRAS expression at the post-transcriptional level, notably through upregulation of microRNAs such as miR-96, which inhibits KRAS mRNA translation (Mazurakova et al. 2023). This indirect regulatory mechanism reduces KRAS protein levels and suppresses KRAS-driven oncogenic signalling pathways, including the downstream PI3K/Akt and MAPK cascades, which are crucial for cancer cell proliferation and survival (Vaidyanathan et al. 2023). Moreover, resveratrol’s polyphenolic structure allows it to bind allosteric or non-canonical sites on KRAS, potentially modulating its function or interfacing with protein-protein interactions without being a classic “drug target” (Saud et al. 2014). This multitarget binding ability is consistent with resveratrol’s broad-spectrum actions affecting numerous cancer-related proteins and pathways simultaneously. Importantly, differential mRNA expression analysis revealed that all hub genes were significantly upregulated in tumor samples, corroborating their relevance in pancreatic cancer progression (Wang et al. 2020). Together, these findings suggest that resveratrol may exert its therapeutic effects through simultaneous modulation of KRAS-driven oncogenesis, EGFR-mediated proliferative signaling, and mTOR-dependent metabolic regulation, thereby targeting the core vulnerabilities of pancreatic cancer biology (Jiang et al. 2009, Xue et al. 2020, Xiao et al. 2021).

## Conclusion

This study established a naturally occurring stilbene polyphenol compound Resveratrol (RVT) screened out most pharmacokinetically favourable on the basis of Lipinski and Veber rule and showed no toxicity predicted through ProTox-3.0 web tool. Resveratrol (RVT) has several target genes against pancreatic cancer like EGFR, SRC, BCL2L1, PTGS2 PI3KCA, PI3KCB and MTOR. These genes are the major hub genes. These hub genes are mainly involved in prime KEGG pathways related to pancreatic cancer. As we know that these hub genes were need to be suppressed as they induce tumour growth, migration, and invasion by different pathways. Our findings on resveratrol indicate that it can effectively influence pancreatic tumor progression by suppressing proliferative and survival signaling, particularly through strong interactions within the PI3K/Akt/mTOR, ERbB and MAPK cascades. Also, promotes apoptosis and overcomes drug resistance by inhibiting survival signaling and altering the tumor microenvironment Moreover, molecular docking results revealed that these hub genes showed the strong binding affinity with Resveratrol. Besides, resveratrol also exhibited strong binding affinity with KRAS, which is not among its predicted target genes, indicating a potential additional mechanism of action. It is important to note that the docking analysis is employed to benchmark specificity, uncover unforeseen interactions, and guide the generation of hypotheses for future research, despite the fact that these genes are not the direct or classic targets of resveratrol. Overall, network pharmacology study based anti-pancreatic cancer potential of resveratrol demonstrates that the compound is a multi-targeted, pharmacokinetically favourable, and synthetically accessible compound with promising therapeutic potential against pancreatic cancer. Its interaction with critical oncogenic pathways and hub proteins highlights its candidacy for further in vivo and clinical validation as a natural anticancer agent. Collectively, these findings provide mechanistic insights into how resveratrol exerts its pleiotropic anticancer activities and support its potential as a multi-targeted therapeutic candidate for pancreatic cancer, either as a stand-alone agent or in combination with conventional treatments.

## CRediT authorship contribution statement

**AB**, Writing-original draft, formal analysis, Data curation, Data analysis, RS Writing-original draft Auto Dock analysis, VJ Data analysis review & editing, SM Proofreading the manuscript. VKS Proofreading the manuscript, Visualisation Data analysis, Data Curation, MKM Proofreading the manuscript, review & editing.

## Acknowledgement

The authors are grateful for the excellent cooperation provided by the Chairman and Director of SR Institute of Management in Lucknow, to conduct this research.

## Figure legends

**Supplementary File 1: Arrangement of common targets of Resveratrol and Pancreatic Cancer on the basis of node degree**. A total 39 common targets which are arranged from maximum node degree to minimum node degree, used to create a Protein-Protein Interaction (PPI) through STRING database.

**Supplementary File 2: Identified hub nodes using the CytoHubba plugin in Cytoscape based on betweenness centrality, node degree, and closeness**. Top 7 in network String0.700_7 july.csv ranked by degree method, including EGFR, SRC, BCL2L1, PTGS2 PI3KCA, PI3KCB and MTOR, were identified as hub nodes.

**Supplementary File 3: Kegg pathway analysis of identified Hub proteins**. Analysis of Involvement of top 7 Hub proteins in different Kegg Pathway on the basis of Enrichment FDR, Fold Enrichment and number of Genes.

Supplementary Figure 1: Core oncogenic interaction network in pancreatic cancer. The network illustrates the strong protein–protein interactions among three key driver genes—**KRAS, TP53**, and **CDKN2A**—which collectively regulate cell proliferation, apoptosis, and cell cycle control. These central molecular nodes represent the major mutational hotspots and signaling hubs contributing to pancreatic tumor initiation and progression.

**Supplementary Figure 2a: Dendrogram representation of enriched KEGG pathways associated with resveratrol-targeted genes in pancreatic cancer**. This hierarchical clustering dendrogram illustrates the interrelationship among significantly enriched pathways obtained from the KEGG database. Pathways closely associated with pancreatic cancer (hsa05212), including PI3K–Akt signaling, ErbB signaling, EGFR tyrosine kinase inhibitor resistance, JAK–STAT signaling, and HIF-1 signaling, are highlighted as key nodes in tumor growth, metabolic regulation, and drug resistance mechanisms. The clustering pattern indicates that resveratrol exerts its anti-pancreatic cancer effects through multi-pathway modulation, primarily targeting oncogenic signaling and metabolic adaptation processes.

**Supplementary Figure 2b**: KEGG pathway–pathway interaction network analysis of overlapping gene targets of resveratrol.

