## Supplementary figures and images for "Deciphering the Therapeutic Potential of Resveratrol Against Pancreatic Cancer Through Network Pharmacology"

### Supplementary Figure 1

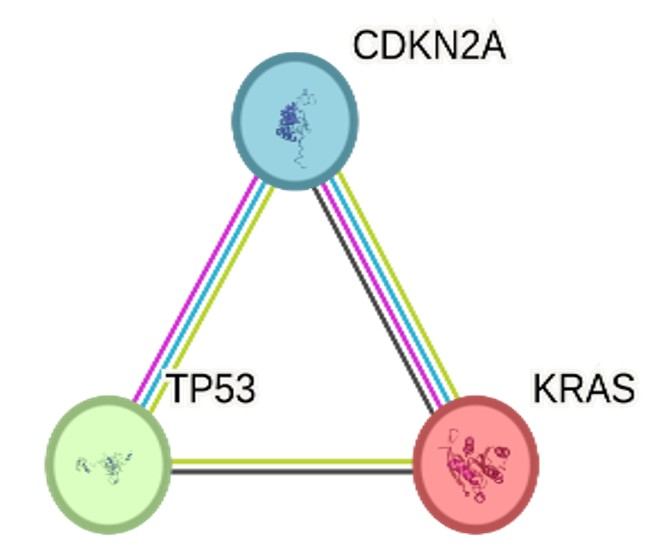

### Supplementary Figure 2

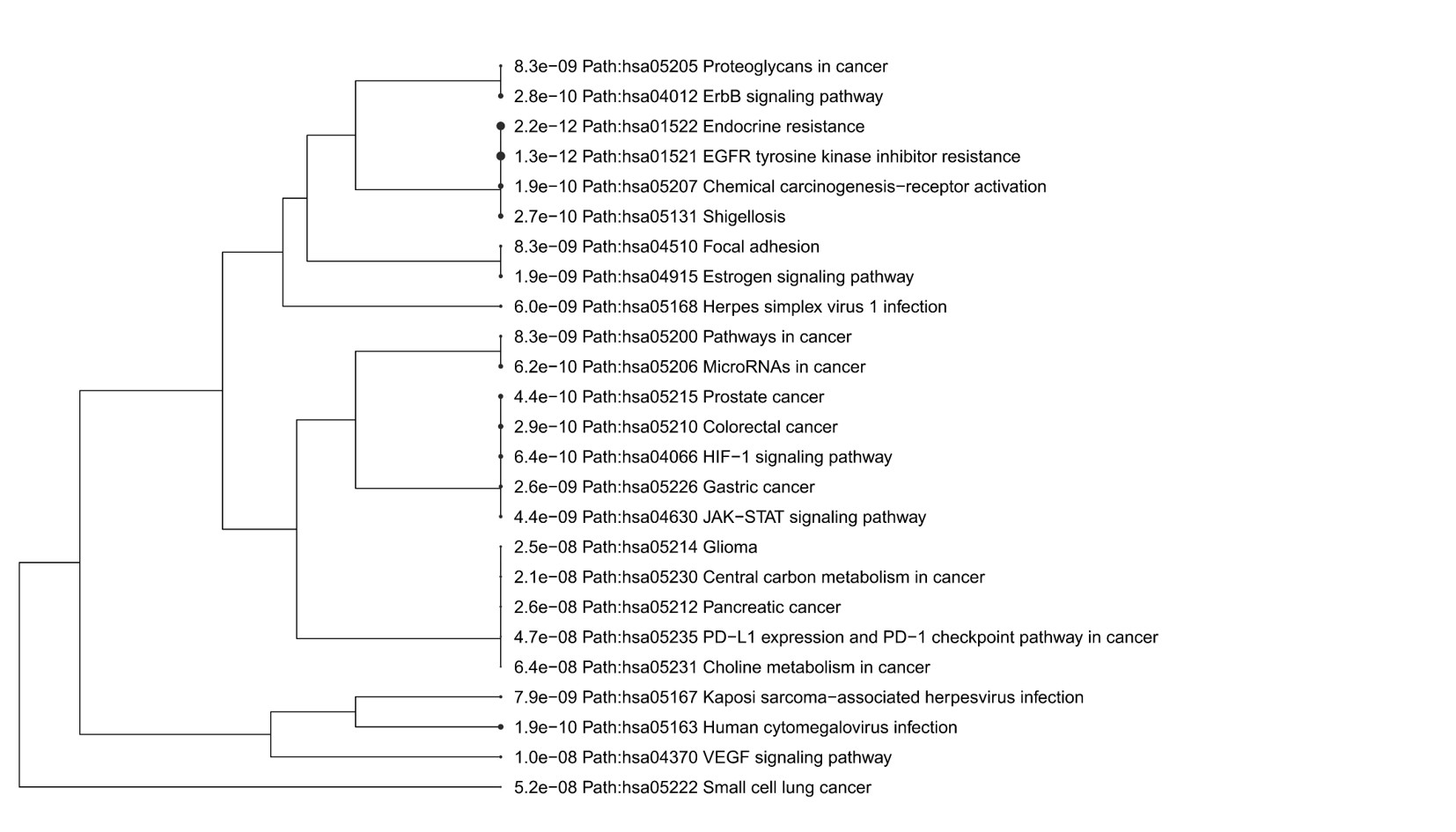

### Supplementary Figure 3

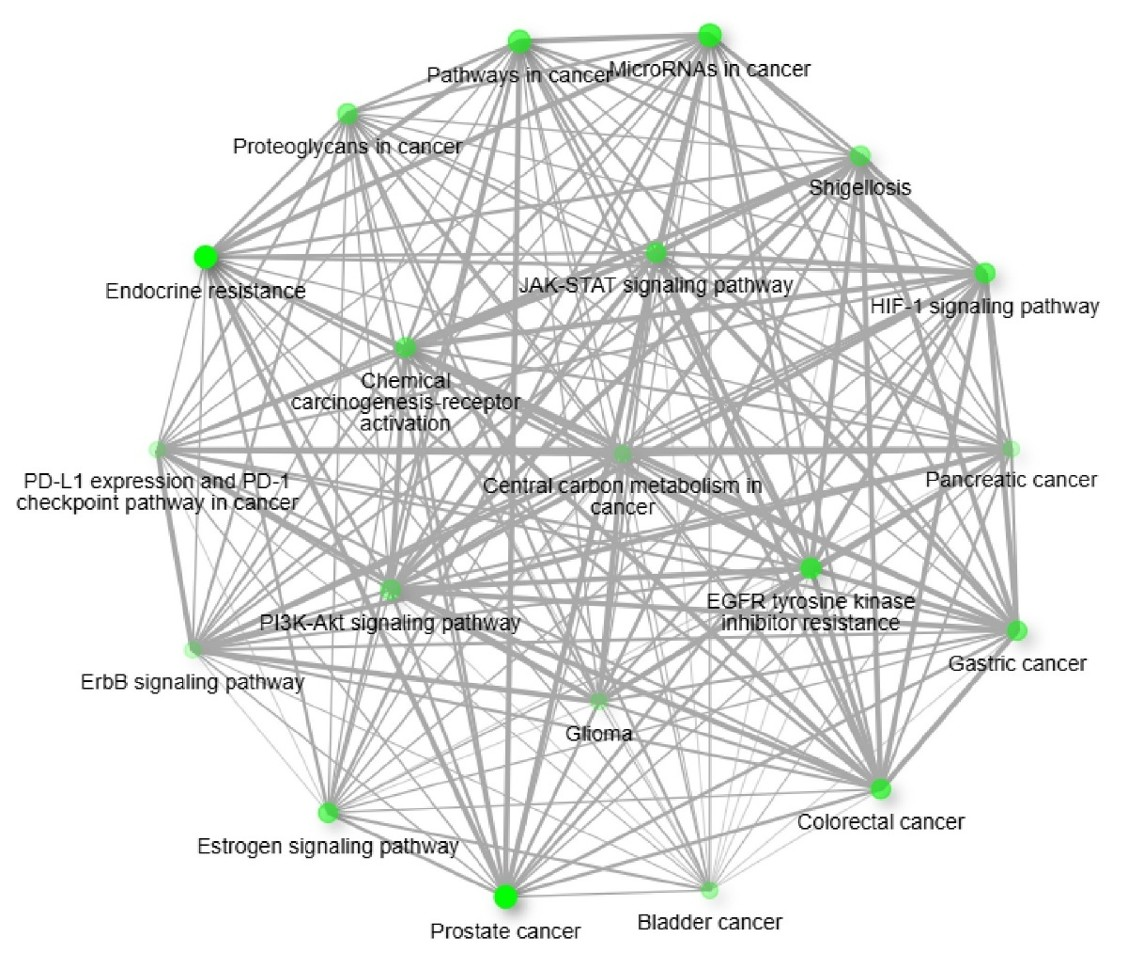
